# Mature tumoroids recapitulate clinically relevant drug response through extended 3D culture in PDAC

**DOI:** 10.64898/2026.04.04.716464

**Authors:** Krzysztof Kuś, David Earnshaw, Artur Piróg, Martyna Siewiera, Sachin Kote, Aleksandra Anna Murzyn, Piotr Świerzewski, Natalia Małek-Trzonkowska, Zuzanna Sandowska-Markiewicz, Katarzyna Unrug-Bielawska, Małgorzata Statkiewicz, Paola Dama, Marcin Przemysław Krzykawski

## Abstract

**Background:** Drug responses in pancreatic ductal adenocarcinoma (PDAC) vary sharply across *in vitro* culture formats, but most 2D–3D comparisons conflate microenvironmental cues with time-dependent cellular adaptation. As a result, conventional assays frequently overestimate drug efficacy and poorly reflect clinical pharmacology.

**Main findings:** We profiled MiaPaCa-2, PANC-1, and CFPAC-1 grown in an extracellular-matrix (ECM) hydrogel for 1–12 days, defining **extended 3D cultures (≥10 days) as *mature tumoroids,*** and quantified 72 h drug responses to a multi-class oncology panel using growth-rate (GR) metrics to normalize for proliferation across formats and durations. Prolonged 3D pre-culture induced broad tolerance, with typical 10–100× reductions in sensitivity to standards of care (5-fluorouracil, SN38, oxaliplatin, gemcitabine, paclitaxel), following a reproducible susceptibility hierarchy (MiaPaCa-2 > PANC-1 > CFPAC-1) after GR correction. In **mature tumoroids**, GR₅₀ values closely approximated clinically observed plasma exposures (e.g., within <4× for 5-FU and <0.5× for gemcitabine), whereas 2D and short-term organoid assays markedly underestimated resistance, often by >100×, thereby overstating drug activity. Notably, CFPAC-1 exhibited increased sensitivity to SN38 and trametinib under mature-organoid conditions, demonstrating that microenvironmental conditioning can invert responses for selected mechanisms. Transcriptomic profiling revealed coordinated up-regulation of multiple ABC transporters with extended 3D residence, tracking resistance phenotypes across lines and implicating transporter-linked tolerance programs.

**Significance:** Together, these data identify **time-in-3D and the emergence of mature tumoroids** as dominant, previously under-controlled determinants of PDAC pharmacology that both induce tolerance and unmask context-dependent vulnerabilities. We propose incorporating both short-term and mature-tumoroid screening arms into preclinical workflows, reporting pre-culture duration alongside GR-normalized effect sizes, and leveraging transporter-informed biomarkers to guide regimen prioritization and sequencing. This framework enhances physiological relevance, reproducibility, and translational fidelity in PDAC drug discovery.

## Introduction

Despite decades of intensive efforts across academia and the pharmaceutical industry, pancreatic ductal adenocarcinoma (PDAC) remains among the most lethal malignancies [1]. Its profound resistance to systemic therapy reflects a convergence of factors, including a dense desmoplastic stroma, aberrant tissue mechanics, restricted drug penetration, immune evasion within the tumor microenvironment, and intrinsic cellular resistance mechanisms [2–4]. Standard chemotherapies provide only marginal benefit: monotherapy with 5-fluorouracil (5-FU) yields a 1-year survival rate of only ∼2%, while gemcitabine modestly improves this to ∼18% [5]. Neither agent is curative; instead, both confer limited survival gains at the cost of substantial toxicity and reduced patient quality of life [5,6].

This persistent therapeutic failure underscores the need for preclinical model systems that more faithfully recapitulate PDAC biology. Three-dimensional (3D) culture systems capture key in vivo hallmarks including cell–cell and cell–matrix interactions, biochemical and oxygen gradients, and altered biomechanics, that remain incompletely represented in traditional two-dimensional (2D) monolayers [7–10]. Within this landscape, engineered 3D systems based on defined matrices offer experimentally tractable platform with a precise control over stiffness, nutrient diffusion, and extracellular-matrix composition [11–13]. In contrast, patient-derived organoids (PDOs) preserve the genomic and phenotypic diversity of human tumors but often rely on poorly defined matrices, complexed and inhibitory cell culture media and display substantial inter-line variability, complicating quantitative drug-response analyses [14–16]. Thus, PDOs are invaluable for capturing patient heterogeneity, whereas engineered 3D systems offer a standardized framework to dissect the microenvironmental determinants of therapeutic response [17,18] [Supplementary Tables S1, S2 and Supplementary Figure S1].

In conventional 2D monolayer cultures, both 5-fluorouracil (5-FU) and gemcitabine display pronounced antiproliferative activity at low micromolar concentrations, indicative of strong in vitro efficacy [19]. Concordantly, patient-derived organoid (PDO) studies, such as those by Tiriac et al. [15] report comparable sensitivity profiles, with robust growth inhibition following exposure to these agents. Together, these findings indicate that both traditional 2D cultures and PDOs capture high apparent susceptibility to these agents under simplified in vitro conditions in comparison to patient serum concentration.

Critically, however, achieving therapeutically relevant effects in vivo requires drug concentrations on the order ∼100-fold higher for 5-FU and ∼700-fold higher for gemcitabine [20–24], levels associated with severe systemic toxicity and limited clinical benefit. This disparity highlights a persistent disconnect between preclinical drug sensitivity and clinical efficacy in PDAC, reinforcing the need for models that better reflect the tumor microenvironment and pharmacologic constraints encountered in patients. Collectively, prior studies highlight a systematic mismatch between model-dependent drug sensitivity and clinical response across PDAC systems, particularly for transporter-linked agents [Table 1]. Notably, this mismatch arises in part because conventional 2D and short-term 3D assays substantially underestimate resistance, yielding effective concentrations far below clinically observed plasma exposures.

**Table 1.** Comparative analysis of drug transporter expression and concordance with clinical drug response across PDAC model systems. This table integrates published evidence with data from the present study to compare expression patterns and functional relevance of selected drug transporters implicated in pancreatic ductal adenocarcinoma (PDAC) chemoresistance across 2D monolayer cultures, extended 3D models, patient-derived organoids (PDOs), and clinical datasets. ABCC3 (MRP3) and ABCA8 are consistently induced following prolonged 3D microenvironmental conditioning and display drug-tolerance associations that align with clinical observations, whereas their expression is absent or variable in 2D cultures and Matrigel-based PDOs. In contrast, ABCB1 (P-gp), ABCG2 (BCRP), and hENT1 (SLC29A1) show context-dependent or inconsistent relationships between in vitro behavior and clinical outcomes, underscoring limitations of conventional models in capturing microenvironment-driven drug transport mechanisms. The consistency verdict indicates the extent to which in vitro transporter behavior recapitulates reported clinical associations with therapeutic response in PDAC (High 3D ↔ clinic, alignment observed only after 3D conditioning; 2D-biased, signal predominant in monolayers but not clinically reproducible; Context-dependent, variable across studies). Abbreviations: ECM, extracellular matrix; PDO, patient-derived organoid; PDX, patient-derived xenograft; KPC, Kras^G12D/+^; Trp53^R172H/+^; Pdx1-Cre mouse model.

| Transporter | 2D cell lines | Extended-3D ("mature tumoroids") | PDOs (Matrigel) | Clinical/Translational evidence | Consistency verdict |
| --- | --- | --- | --- | --- | --- |
| **ABCC3 (MRP3)** | Typically low or baseline unless selected by drug exposure | Strongly induced after prolonged 3D residence ($\geq 8$ days) **in our models**; associated with hypoxia and ECM-stress signatures | Variable; affected by Matrigel lot and differentiation state | Overexpressed in tumors; correlates with chemoresistance; inhibition (MCI-715) restores sensitivity in xenograft, PDX and KPC models [31] | **High 3D <-> clinic, not obvious in 2D/PDO ->** Strong candidate |
| **ABCA8** | Minimal or inconsistent | Induced under bile-acid-rich or hypoxic 3D conditions **in selected PDAC models**; linked to S1PR2-ERK activation and gemcitabine tolerance. | Variable expression; unstable across passages | Clinical data **from recent cohorts** suggest an ABCA8-S1PR2-ERK axis correlates with poor gemcitabine response and reduced survival [34] | **High 3D <-> clinic**, frequently under-captured in 2D/PDO systems -> Strong candidate |
| **ABCB1 (P-gp)** | Commonly high after drug selection | Not specifically induced by 3D residence | Mixed; moderate to high expression in some lines | Inconsistent prognostic value; some studies link high ABCB1 to poor FOLFIRINOX outcome, others neutral [96] | **2D-biased signal**; clinic **mixed** -> limited translational alignment |
| **ABCG2 (BCRP)** | Frequently elevated in selected or stem-like subclones | No reproducible 3D-specific induction | Mixed; dependent on culture stress | Newer cohort suggests ABCG2/ABCB1 expression may impact Gem/Nab outcomes, but results remain variable across studies [97] | **Context-dependent**, clinic mixed -> Lower priority |
| **hENT1 (SLC29A1) (uptake)** | Robust predictor in controlled gemcitabine sensitivity in vitro | Often down-regulated under TGF-beta or hypoxic 3D conditions | Variable staining and assay-dependent expression | Adjuvant trials: positive predictive signal; advanced disease/cohort show **conflicting** results; assay standardization lacking [98] | **2D overestimates**; 3D may better reflect clinical down-modulation, but heterogeneity limits clarity |

Multiple, non-exclusive mechanisms contribute to the reduced drug sensitivity observed in 3D compared with 2D cultures, including physical barriers to drug diffusion imposed by tissue density, cell adhesion, and extracellular-matrix (ECM) deposition [25];hypoxia-driven induction of efflux transporters such as ABCB1 (MDR1) [26,27]; and broad metabolic adaptations encompassing quiescence, altered redox and pH states, and nutrient limitation. Together, these processes reshape cellular metabolism and impair the efficacy of replication-dependent agents [28,29].

Beyond ABCB1, PDAC exhibits widespread dysregulation of ATP-binding cassette (ABC) transporters. Tumor profiling studies report consistent up-regulation of ABCC and ABCG family members, alongside altered cholesterol-handling transporters largely independent of KRAS exon 2 status [30]. Functionally, ABCC3 is recurrently elevated in PDAC [31] and has been linked to decreased sensitivity to multiple chemotherapeutics, including epipodophyllotoxins and antimetabolites, with clinical data suggesting a contribution to poor treatment response [32,33]. In parallel, ABCA8 has been implicated in gemcitabine insensitivity through a taurocholic-acid/S1PR2-ERK signaling axis, highlighting transporter functions that promote drug resistance via pathway activation rather than physical drug extrusion [34]. Consistent with these observations, PDAC cell-line models resistant to 5-FU or gemcitabine exhibit up-regulation of overlapping ABCC family members, with functional evidence implicating ABCC5 in both contexts [35,36]. The commonly used PDAC cell lines employed in this study harbor canonical driver mutations representative of the disease, including alterations in KRAS, TP53, CDKN2A, and SMAD4 [Supplementary Figure S2].

Collectively, transporter and matrix-driven programs foster multidrug-tolerant phenotypes, aligning with the broader “pan-resistance” framework described in PDAC and other solid tumors [37,38]. However, most prior studies compare rapidly dividing 2D monolayers with 3D spheroids of heterogeneous and often poorly defined culture durations, thereby conflating culture format with time-dependent cellular adaptation variable known to influence diffusion barriers, hypoxia, and cell-cycle state [39,40]. In this context, we define “mature tumoroids” as tumor 3DCC (3D cell culture) that have undergone prolonged residence in a defined ECM-containing 3D culture, permitting time-dependent microenvironmental conditioning, including structural compaction, diffusion and oxygen gradients, and metabolic and transcriptional remodeling, to emerge prior to pharmacologic interrogation. Moreover, conventional IC₅₀-based analyses are highly sensitive to proliferation differences across experimental conditions. Growth-rate (GR)–based metrics address this limitation by normalizing drug response to division rate, thereby enabling equitable comparisons across culture formats and temporal states [41].

Notably, CFPAC-1 cells displayed increased sensitivity in 3D to selected agents, including SN38 and trametinib, demonstrating that microenvironmental conditioning can invert drug responses in a mechanism-dependent manner. Consistent with transporter-linked tolerance, transcriptomic profiling revealed induction of resistance-associated programs, including ABCC3 and ABCA8, that tracked with drug-response patterns across cell lines [31,34]. In MiaPaCa-2 cells, prolonged residence in 3D culture was associated with the emergence of a broad drug-tolerant phenotype accompanied by increased expression of ABCA-family transporters, with tolerance magnitude varying by agent, underscoring the non-uniform and mechanism-dependent nature of 3D-induced resistance [42,43].

Together, these findings underscore that most conventional 2D–3D comparisons confound culture format with adaptation time, as 3DCC “age” alone reshapes diffusion barriers and toxicity profiles, while community standards continue to highlight heterogeneous and frequently under-reported 3D culture parameters [39,44].

By explicitly varying time-in-3D and applying GR-normalized analytics, we avoid proliferation confounders inherent to IC50/EC50-based analyses and enable fair cross-condition comparisons [45]. Our findings demonstrate that prolonged 3D residence is a dominant driver of broad tolerance and context-specific vulnerabilities, consistent with the heightened chemoresistance observed in matrix-rich, stroma-like PDAC spheroids [42].

Translationally, extended 3D conditioning more closely recapitulates chronic stromal exposure and treatment history in vivo, consistent with the objectives of New Approach Methodologies (NAMs) to improve the physiological relevance and predictive performance of human-based preclinical models. In this context, prolonged 3D culture can uncover context-dependent therapeutic liabilities, such as MEK-pathway sensitivity, that are not apparent in short-term assays or rapidly proliferating 2D systems [46–48].

We therefore propose incorporating paired short- and extended-3D arms into preclinical screening workflows as a pragmatic NAM-aligned strategy to prioritize regimens active against pre-conditioned tumors, identify transporter-linked biomarkers for stratification, and inform rational sequencing and combination strategies. Routine reporting of 3D pre-culture duration alongside growth-rate (GR)–normalized effect sizes would further align with emerging regulatory expectations for transparency, reproducibility, and cross-study comparability in PDAC drug-discovery and translational research pipelines [45,49].

## Results

A central concern in PDAC pharmacology is that most 2D–3D comparisons conflate microenvironmental format with time-in-3D (i.e., adaptation or “age”), obscuring whether observed drug-response differences reflect matrix composition or progressive cellular conditioning. We therefore asked whether residence time in a *defined* 3D ECM is itself a driver of drug response, and if so, whether it (i) broadly reduces sensitivity, (ii) can invert sensitivity for specific drug mechanisms, (iii) follows a reproducible susceptibility ranking by cell line, and (iv) correlates with transporter programs that could mechanistically explain the observed functional changes. To avoid proliferation bias across formats and temporal states, all dose–responses were analyzed using growth rate–normalized (GR) metrics, enabling direct comparison of cytostatic and cytotoxic effects independent of baseline growth differences. Complete GR curves are shown in Figures 1–3, while quantitative response parameters (IC₅₀, GR₅₀, GR_AOC) across formats and timepoints are reported in Tables S3–S5.

**Figure 1.**
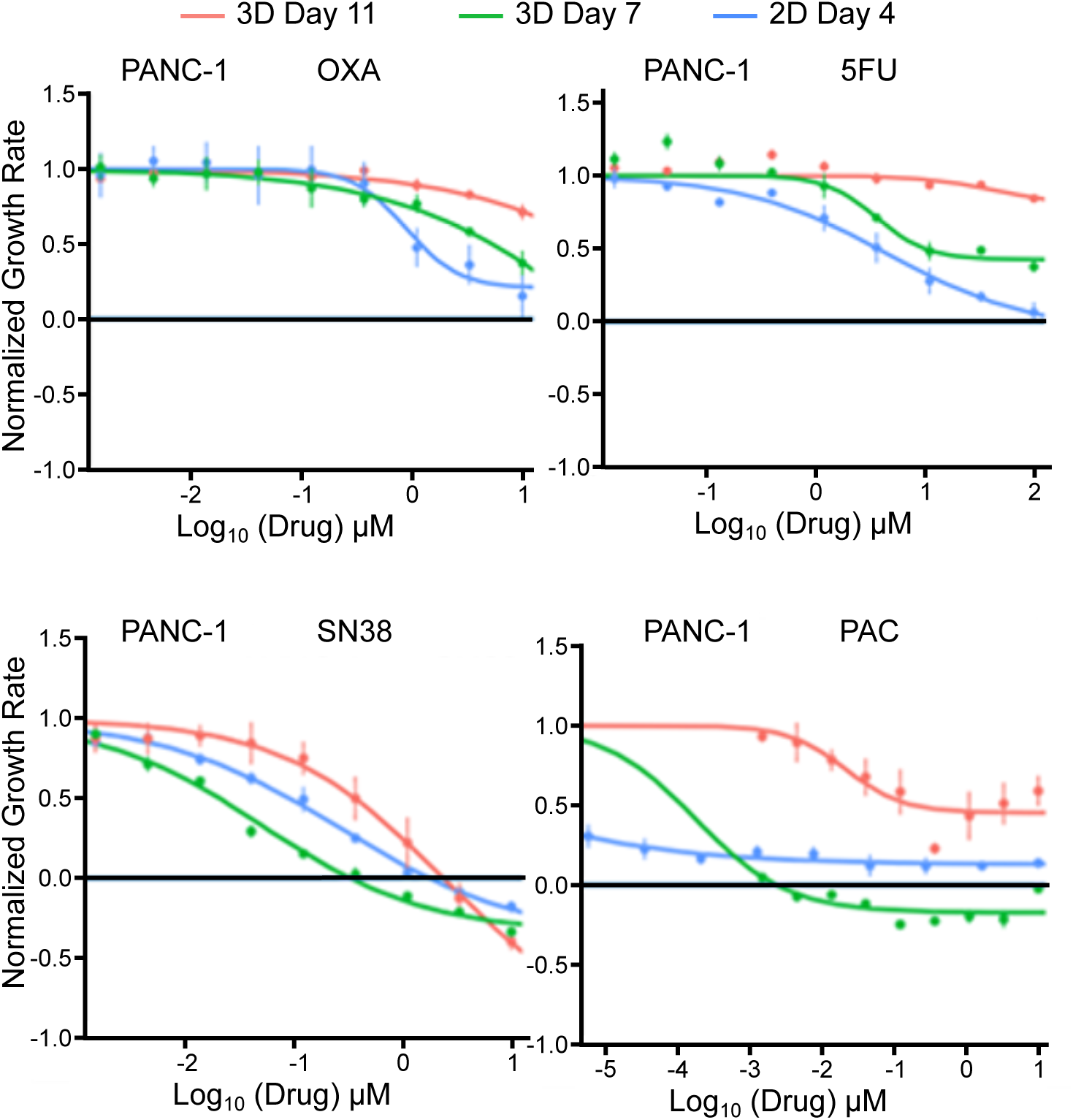
Time-in-3D pre-conditioning reduces drug sensitivity in PANC-1 spheroids. Growth-rate–normalized dose–response (GR) curves for PANC-1 cultured in an ECM-containing 3D hydrogel and with 4 different anti-cancer drugs for 72h. Upper panels: oxaliplatin (OXA, left) and 5-fluorouracil (5FU, right) comparing 2D day-4 cultures (blue) with 3D pre-conditioned cultures at day 7, 11. Bottom panels: SN38 (left) and paclitaxel (PAC, right) comparing 2D day-4 (blue) with 3D day-7 (green) and 3D day-11 (red). Points show mean ± SEM.; horizontal black line marks GR = 0 (cytostatic threshold). Axes: x, log_10_ (drug) (µM); y, normalized GR.

**Figure 2.**
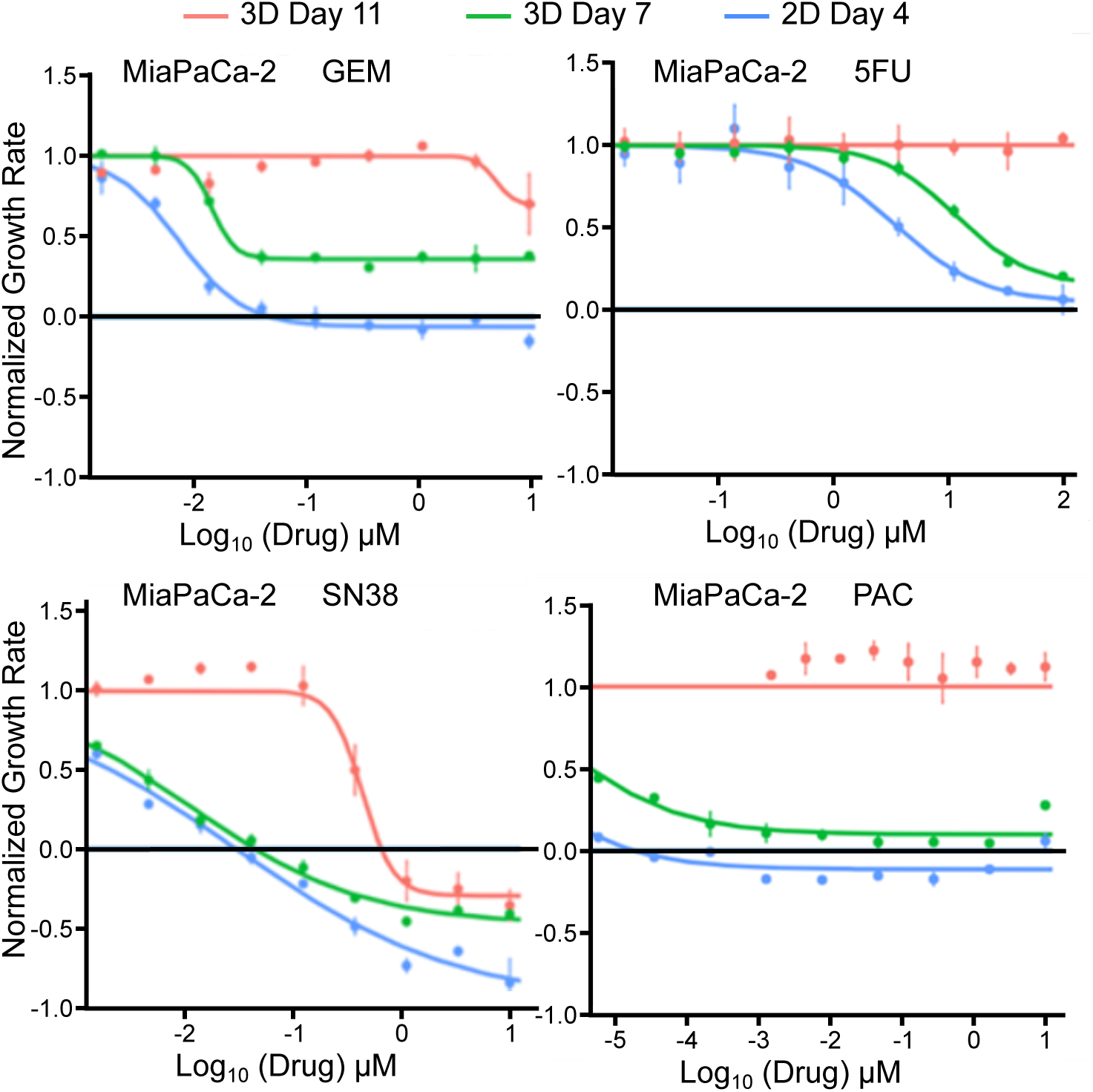
Time-in-3D pre-conditioning reduces drug sensitivity in MiaPaCa-2 spheroids. Growth-rate–normalized dose–response (GR) curves for MiaPaCa-2 spheroids pre-cultured in an ECM-containing 3D hydrogel for 4, 7, 11 days, then treated for 72 h with (upper panels) gemcitabine (GEM, left) and 5-fluorouracil (5FU, right) or (bottom panels) SN38 (left) and paclitaxel (PAC, right). Points show mean ± SEM.; horizontal black line marks GR = 0 (cytostatic threshold). Axes: x, log_10_ (drug) (µM); y, normalized GR.

**Figure 3.**
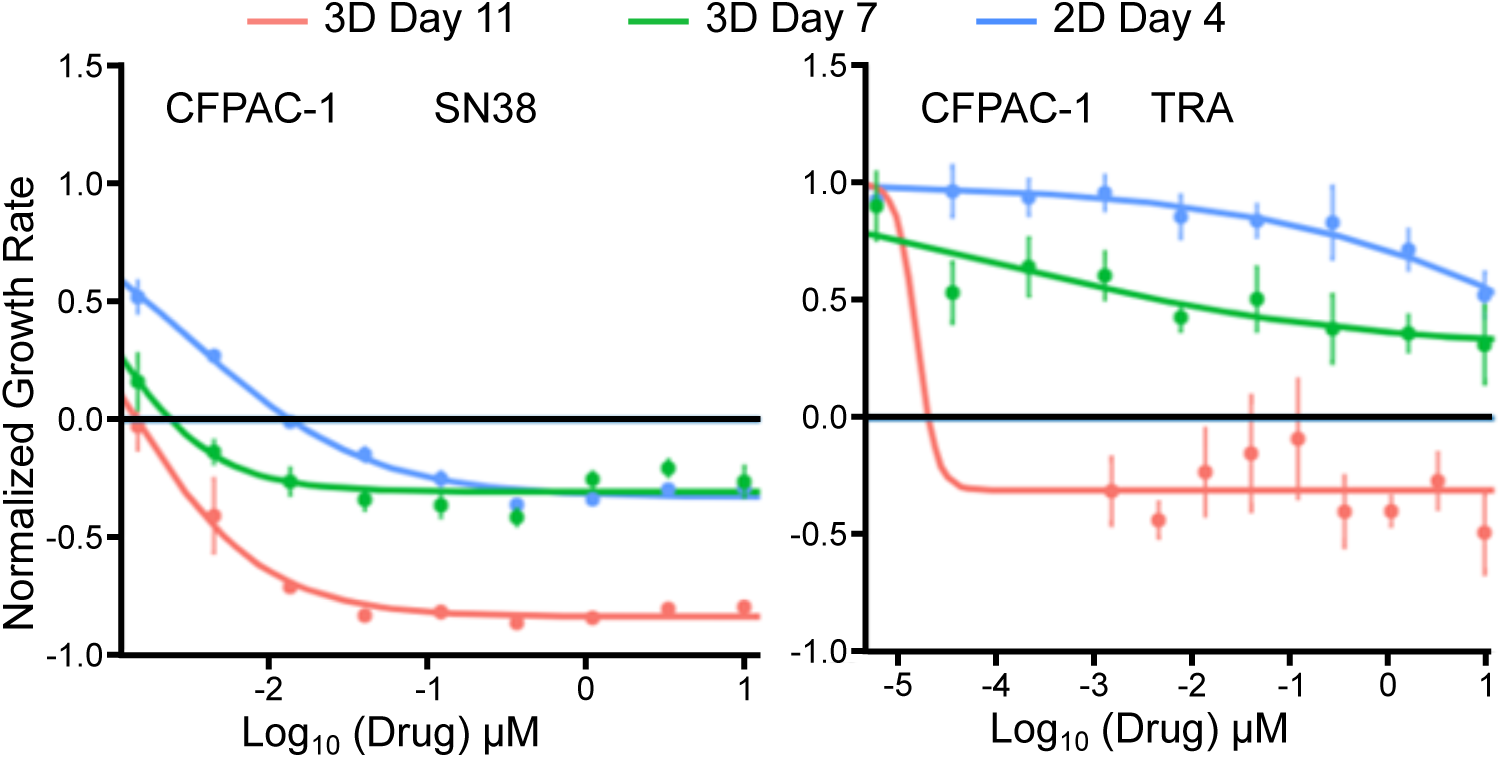
Time-in-3D pre-conditioning can increase drug sensitivity in CFPAC-1. Growth rate–normalized dose–response (GR) curves for CFPAC-1 cultured either in 2D for 4 days (blue) or pre-cultured in an ECM-containing 3D hydrogel for 7 days (green) or 11 days (red), then treated for 72 h with upper panel: SN38 or bottom panel: trametinib (TRA). Points represent mean ± SEM.; the horizontal black line indicates GR = 0 (cytostatic threshold). Axes: x, log_10_ (drug) (μM); y, normalized GR.

### Time-in-3D pre-conditioning reduces drug sensitivity in PANC-1 and MiaPaCa2

Across repeated, GR-normalized 72-h dose–response assays, increasing residence time in the 3D ECM hydrogel produced systematic reduced potency and reduced maximal effect in both cell lines. In PANC-1, oxaliplatin and 5-fluorouracil showed the most prominent rightward shifts, while SN38 and paclitaxel displayed attenuation of both potency and depth of kill by day 11 (Figure 1). In MiaPaCa-2, gemcitabine, 5-fluorouracil, SN38, and paclitaxel all moved toward lower apparent activity with 3DCC age, with gemcitabine showing one of the steepest losses between days 7 and 11 (Figure 2). Consistent with these curves shifts, activity tallies declined with time: PANC-1 retained 5/5 SOC (14/14 total) at day 7, decreasing to 2/5 (11/14 total) at day 11 (Supplementary Table S3), whereas MiaPaCa-2 retained activity for 3/5 SOC (9/14 total agents) at 3D day 7 but only 0/5 (2/14 total) by day 11 (Supplementary Table S4).

Two features of the GR curves clarify the underlying biology. First, potency loss precedes efficacy loss for several agents: the half-maximal inhibition (GR50) values shifted to higher concentrations before the high-dose plateau (GRmax) rebounded toward zero inhibition, suggesting that early resistance primarily reflects a higher exposure requirement rather than complete loss of the druggable mechanism. Second, Hill-slope shallowing at day 11, visible for select drug–line pairs, indicates increased response heterogeneity, consistent with microenvironment-driven phenotypic diversification within aging spheroids.

Importantly, these time-dependent changes were reproducible across biological replicates and persisted after controlling for well position and batch, indicating that they are not technical artifacts. Parallel growth tracking in 3D showed continued mass expansion into the second week (Supplementary Figure S3), highlighting the importance of GR normalization to distinguish true resistance from proliferation-driven effects. Together, the curve geometry, the activity tallies, and the replicate stability indicate that extended residence in a stromal-like 3D context pushes PANC-1 and MiaPaCa-2 toward multidrug tolerance, with the magnitude and kinetics of this shift varying by drug class and cell line.

### Extended 3D conditioning shifts in-vitro effective concentrations toward clinically relevant exposure ranges

To benchmark these in vitro resistance shifts against physiologically relevant exposures, we compared GR₅₀ values from 2D and Day-11 3D cultures with reported mean patient plasma Cmax levels for the same standard-of-care agents (Supplementary Table S6). Whereas Cmax values exceeded 2D GR₅₀s by one to two orders of magnitude, they were comparable to, or in several cases lower than, the Day-11 3D GR₅₀s, particularly for MiaPaCa-2. For instance, 5-fluorouracil, gemcitabine, and paclitaxel each exhibited >10-fold GR₅₀ elevations in Day-11 3D MiaPaCa-2 spheroids, bringing in-vitro effective concentrations into the range of clinically observed plasma Cmax values. Similar but less pronounced patterns were observed in PANC-1, whereas CFPAC-1 remained largely below clinical exposure thresholds for standard-of-care agents (Figure 3; Supplementary Table S6). These data confirm that extended 3D residence not only reduces apparent potency but also repositions in vitro drug response into a clinically relevant concentration window, underscoring the translational relevance of time-dependent 3D conditioning.

### Functional tolerance aligns with transporter-associated transcriptional remodeling

Finally, these functional trends align with the transcriptomic patterns observed conditions exhibiting the largest ΔGR_AOC losses tend to coincide with up-regulation of ABC transporters at day 11, coinciding with increased expression of subset of drug resistance genes and ABC transporters at day 11 (Figure 4; Figure 5) [50], consistent with diminished intracellular drug exposure and blunted cytotoxic responses after prolonged 3D conditioning. Day-11 curves that exhibit shallower slopes and higher high-dose plateaus further indicate coexistence of more- and less-responsive subpopulations rather than a uniform shift; this functional heterogeneity mirrors the cell line-specific, gene-family-skewed ABC-transporter induction, supporting a model in which microenvironment-driven state diversity, rather than simple penetration limits or growth arrest, underlies the broadened tolerance profile after extended 3D residence.

**Figure 4.**
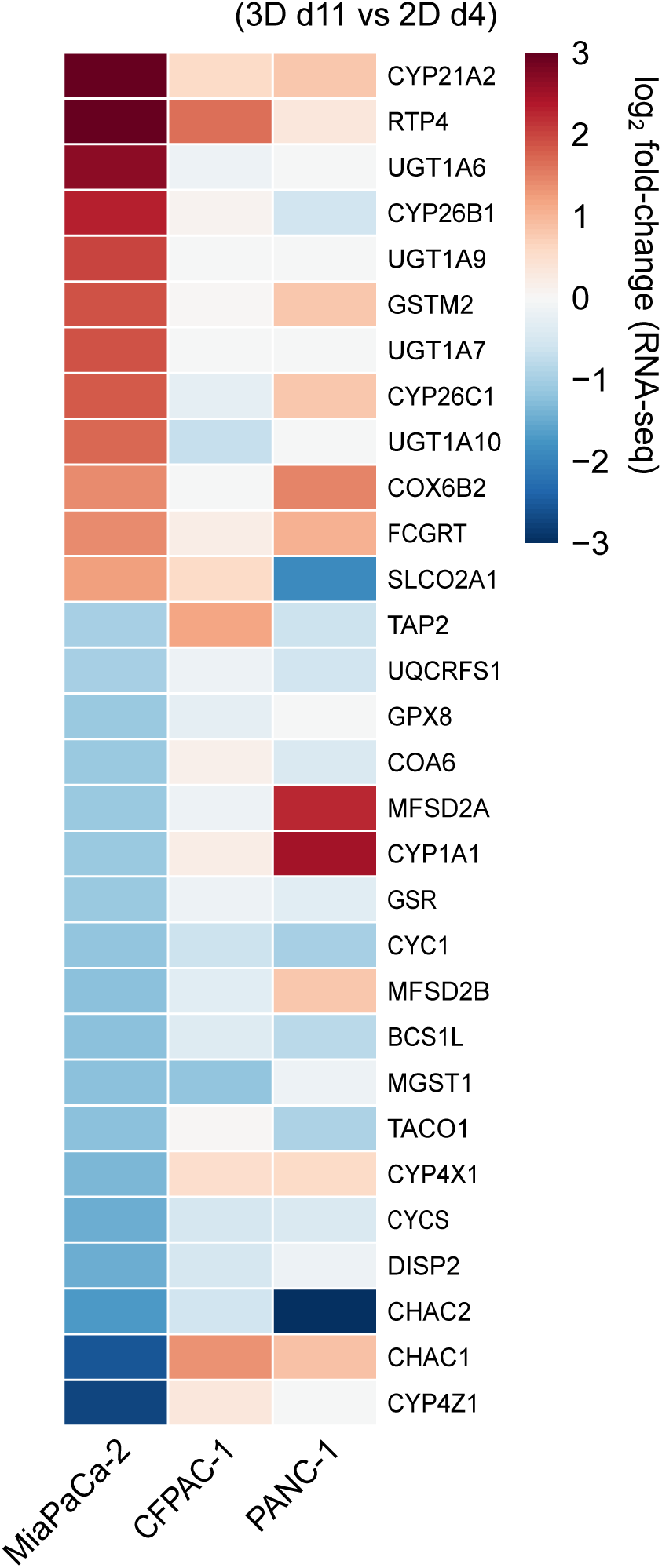
Transcriptional induction of drug resistance genes under prolonged 3D conditioning. Heatmap showing transcriptional log₂ fold-change (3D d11 vs 2D d4) for a selection of drug resistance genes in MiaPaCa-2, CFPAC-1, and PANC-1 cells [50]. Extended 3D culture induces broad upregulation of multiple genes in MiaPaCa-2, whereas CFPAC-1 exhibits attenuated or selective responses. These patterns align with differential GR-based tolerance phenotypes observed across models.

**Figure 5.**
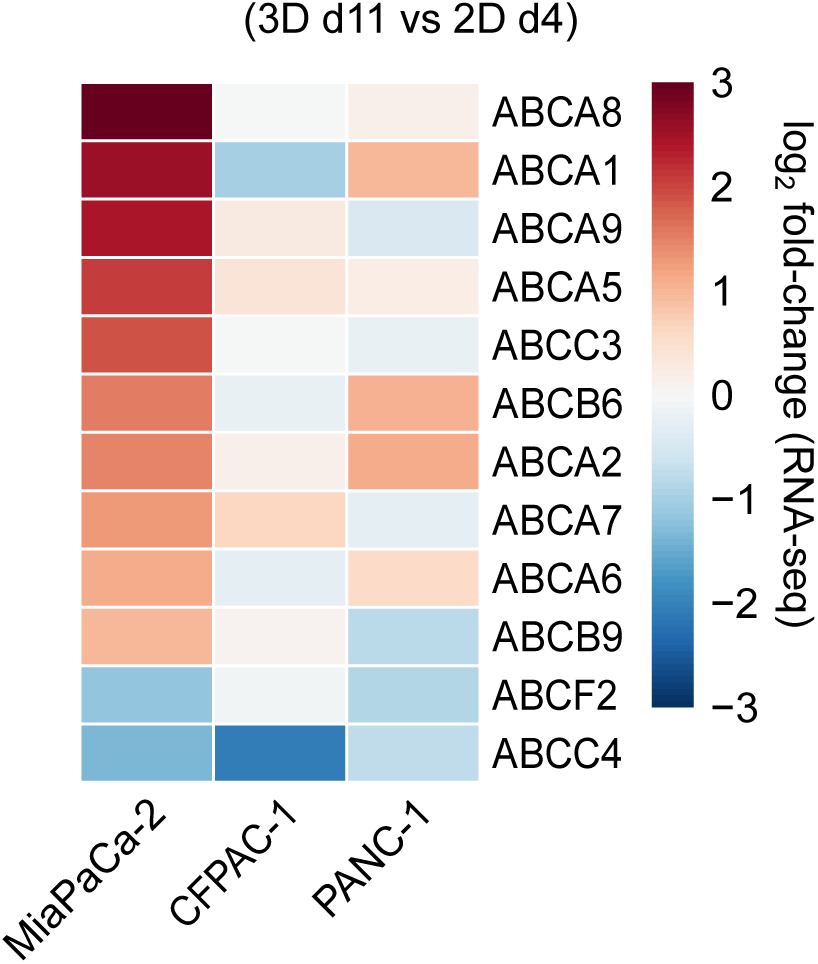
Coordinated induction of ABC transporters under prolonged 3D conditioning. Log₂ fold-change (3D d11 vs 2D d4) heatmap of ATP-binding cassette (ABC) transporter expression in MiaPaCa-2, CFPAC-1, and PANC-1 cells. Extended 3D culture induces broad upregulation of multiple ABCA family members in MiaPaCa-2, while CFPAC-1 shows attenuated induction and PANC-1 exhibits heterogeneous shifts. Downregulation of selected transporters (e.g., ABCF2, ABCC4) underscores the non-uniform nature of 3D-associated transcriptional remodeling. These data support engagement of efflux-associated programs under prolonged 3D residence [50].

### Extended 3D residence unmasks sensitivity inversions in CFPAC-1

In contrast to the global tolerance trend observed in PANC-1 and MiaPaCa-2, CFPAC-1 exhibited increased sensitivity with prolonged residence in 3D for selected drug mechanisms. GR-normalized dose–response curves for SN38 and trametinib shifted downward and leftward as pre-culture progressed from day 7 to day 11, with the largest gains at day 11 reflected by reduced GR50 and GR_AOC values (Figure 3). These shifts were reproducible across replicates and accompanied by consistent curve geometry, indicating a genuine increase in drug efficacy rather than technical variability. Consistent with the curve-level analysis, CFPAC-1 retained activity for 4/5 standard-of-care agents (13/14 total compounds) at both 3D day 7 and day 11 (Supplementary Table S5), with statistically significant changes encoded at the GR_AOC level (p < 0.05; p < 0.01). Notably, SN38 and trametinib met predefined thresholds for increased sensitivity (≥5-fold change in IC₅₀/GR₅₀ or ≥1.5-fold change in GR_AOC), as indicated by green-coded entries. This gain-of-sensitivity phenotype contrasts with the predominantly tolerance-driven responses observed in PANC-1 and MiaPaCa-2 under extended 3D conditioning.

### Sensitivity inversions occur in the absence of transporter upregulation

These findings indicate that prolonged 3D residence can yield qualitatively distinct response trajectories, including sensitivity inversions, rather than uniformly promoting drug tolerance. To explore correlations of these functional inversions, we compared changes in GR_AOC with transporter expression profiles across cell lines and conditions. In concordance, RNA-seq data comparison revealed that sensitivity gains in 3D tend to occur under conditions of minimal or absent ABC-transporter induction, whereas tolerance phenotypes frequently co-occur with increased transporter expression (Figure 5) [50]. This qualitative anti-correlation is consistent with transporter capacity and its upstream regulatory context, modulating whether prolonged 3D residence manifests as tolerance or sensitivity inversion, as observed for SN38 and trametinib in CFPAC-1.

### Rank order of susceptibility across lines

Aggregating GR-normalized response shifts across the full drug panel revealed a reproducible rank order of susceptibility to time-in-3D–induced tolerance: MiaPaCa-2 > PANC-1 > CFPAC-1.

This order is supported by multiple concordant readouts. First, MiaPaCa-2 exhibited pronounced rightward and upward shifts in dose–response curves, accompanied by near-complete loss of standard-of-care activity by day 11 (Figure 2). Consistent with this, MiaPaCa-2 maintained activity for 3/5 standard-of-care agents at 3D day 7 but 0/5 at day 11 (Supplementary Table S4).

Second, PANC-1 displayed broader but intermediate attenuation across the drug panel, with clear reductions in both potency and efficacy emerging by day 11 (Figure 1). Activity tallies declined from 5/5 standard-of-care agents on day 7 to 2/5 at day 11 (Supplementary Table S3), placing PANC-1 between MiaPaCa-2 and CFPAC-1 in susceptibility to extended 3D conditioning.

Finally, CFPAC-1 maintained the broadest spectrum of drug activity under prolonged 3D residence, retaining 4/5 standard-of-care agents at both day 7 and day 11, together with selective sensitivity gains for specific mechanisms (Figure 3; Supplementary Table S5).

Together, these concordant functional metrics establish a cell-line-specific hierarchy of microenvironmental susceptibility, in which extended 3D residence drives rapid and profound tolerance in MiaPaCa-2, intermediate adaptation in PANC-1, and relative stability or sensitivity inversion in CFPAC-1.

### Transcriptional Reprogramming Induced by Prolonged 3D Culture

A curated resistance-gene heatmap comparing 3D day 11 to 2D day 4 revealed a coordinated transcriptomic shift across all three PDAC cell lines (Figure 4). Prolonged residence in 3D was associated with broad upregulation of resistance-linked genes, with functional classes consistently enriched across models. Expression values are reported as log₂ fold-changes, enabling direct comparison of both magnitude and direction across conditions.

Focusing on membrane transport processes, members of the ATP-binding cassette (ABC) superfamily exhibited pronounced induction following extended 3D culture (Figure 5). The ABC transporter heatmap (ABCA/ABCB/ABCC/ABCG), organized by expression similarity, shows a widespread upward shift in transporter expression in 3D day 11 relative to 2D day 4. Regions of strong transporter induction align with conditions exhibiting reduced drug sensitivity in GR-based analyses (ΔGR_AOC), whereas conditions with preserved or increased sensitivity show minimal or attenuated transporter changes. This concordance supports a relationship between ABC transporter activation and tolerance-associated phenotypes, with loss of sensitivity co-localizing with transporter upregulation and sensitivity gains occurring in contexts of limited induction.

Cell line–specific patterns further refine this relationship. In MiaPaCa-2 and PANC-1, widespread transporter upregulation coincides with broad GR-normalized tolerance across drug classes. In contrast, CFPAC-1 displays attenuated transporter induction in conditions associated with increased sensitivity (e.g., SN38 and trametinib), consistent with the observed sensitivity inversion. While these analyses are correlative, the alignment between transcriptional and functional shifts implicates efflux-associated programs as candidate contributors to the pharmacologic phenotypes emerging under prolonged 3D conditioning.

Where assessed at the protein level, transcriptomic changes were directionally supported by proteomic measurements (Figure 7). Notably, ABCA8 exhibited consistent induction at both RNA and protein levels, while ABCB6 showed concordant but more variable protein-level increases. Although proteomic coverage was not uniform across all transporters, these cross-modal observations reinforce the functional relevance of 3D-associated transcriptional reprogramming while remaining within the descriptive scope of the dataset.

### Concordance between functional shifts and transporter expression

To relate functional drug responses to putative resistance programs, we overlaid the ΔGR_AOC map (3D–2D) with matched ABC-transporter expression heatmaps measured at the same timepoints (Figure 6) [50]. A clear and consistent pattern emerged: losses of sensitivity in 3D co-localize with transporter up-regulation, most prominently within the ABCC (MRP) and ABCB family members, whereas gains in 3D sensitivity such as those observed in CFPAC-1 with SN38 and trametinib, aligned with lower or unchanged ABC expression. This inverse relationship was evident both across drugs within a given cell line and across cell lines for the same drug, indicating that it is not driven by a single compound or line-specific idiosyncrasy.

**Figure 6.**
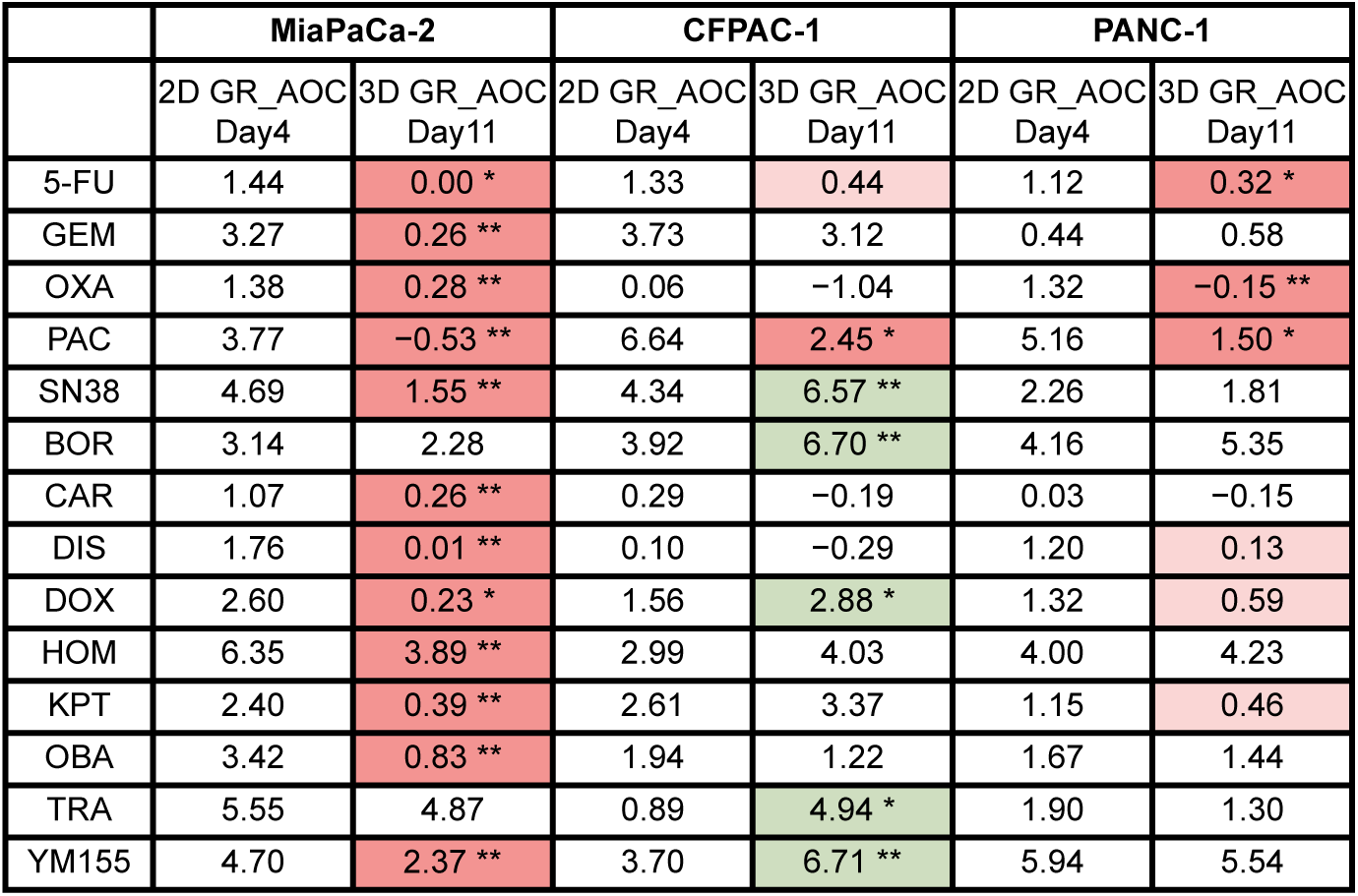
Functional drug-response shifts. Summary of ΔGR_AOC values (3D Day 11 − 2D Day 4; unitless) for each drug–cell-line pair (MiaPaCa-2, PANC-1, CFPAC-1). Boxes indicate the magnitude and significance of activity changes relative to 2D culture (pink, reduced sensitivity in 3D; green, increased sensitivity in 3D; *P<0.05, **P<0.01).

**Figure 7.**
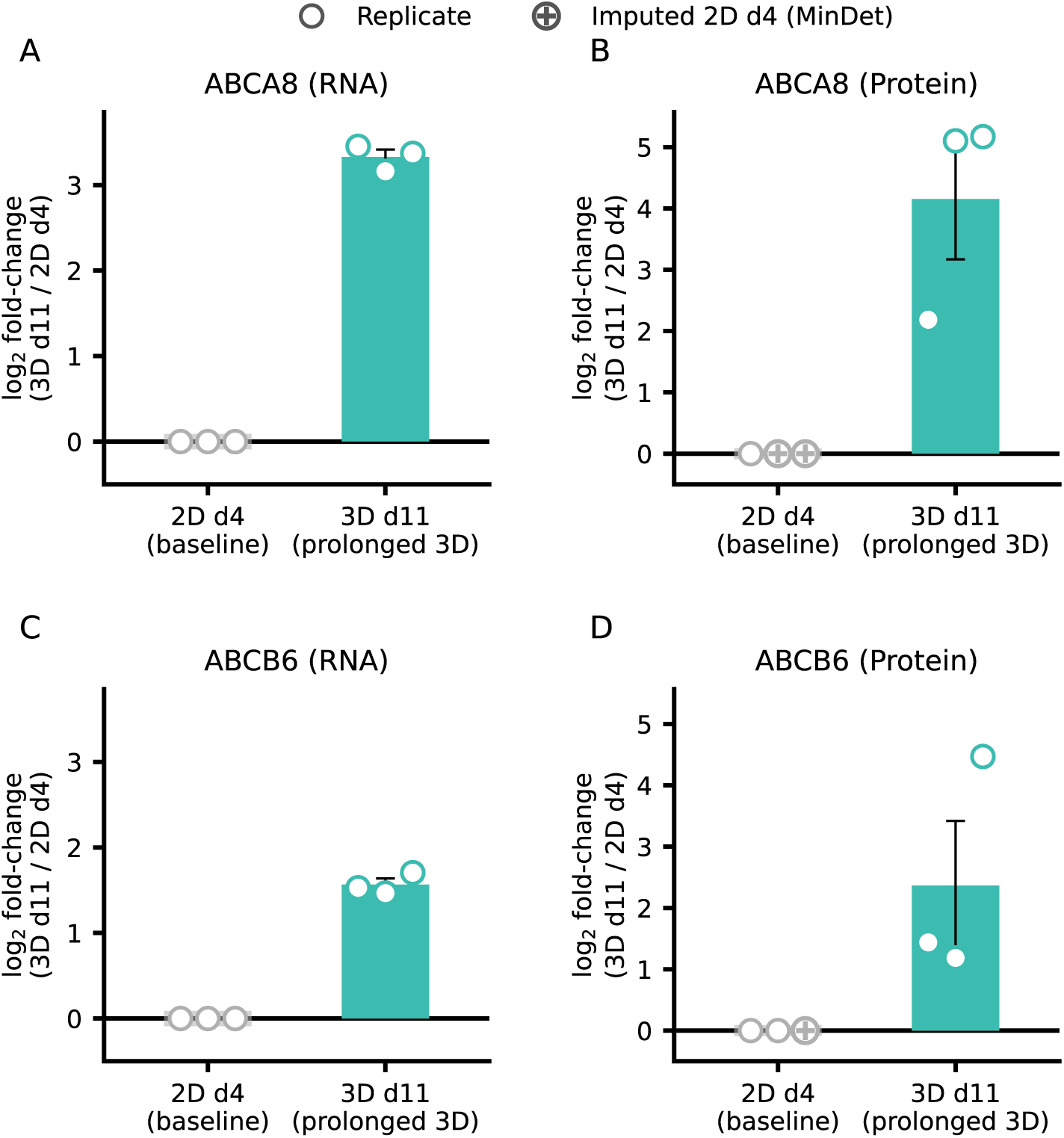
Cross-modal validation of transporter induction in MiaPaCa-2 under prolonged 3D culture. ABCA8 and ABCB6 expression was assessed at the RNA level (A, C) [50] and protein level (B, D) in MiaPaCa-2 cells comparing 2D d4 (baseline) versus 3D d11 (prolonged 3D). Values are shown as log₂ fold-change (3D d11 / 2D d4) computed from paired measurements within each modality. Points represent biological replicates; summary bars indicate mean ± SEM. For proteomics, missing values were imputed using a MinDet strategy as indicated. ABCA8 shows strong directionally concordant induction across RNA and protein, while ABCB6 exhibits concordant induction with increased variability at the protein level.

Two features further strengthened this concordance. First, magnitude coupling drug–line pairs exhibiting the largest positive ΔGR_AOC value (strongest functional tolerance) also tended to show the greatest aggregate ABC induction, quantified as the summed of standardized log2 fold-changes across ABC genes. This pattern is consistent with a graded, dose–response–like association between transporter program intensity and phenotypic tolerance. Second, family specificity: expression shifts were enriched among ABCC (MRP) transporters, with more modest or heterogeneous changes among ABCA lipid-handling transporters. This distribution is consistent with efflux capacity being a principal correlation of reduced intracellular drug exposure, while ABCA-associated changes may contribute indirectly via membrane or lipid remodeling. Importantly, discordant quadrants, drug–line pairs where ΔGR_AOC increased without corresponding ABC induction, or in which ABC expression rose without measurable functional tolerance, were infrequent and line-specific. Notably, in CFPAC-1, the SN38 and trametinib inversions localized to the “gain/low-ABC” quadrant characterized by increased 3D sensitivity in the context of muted ABC transporter induction, consistent with the limited efflux induction permitting the emergence of context-dependent pathway vulnerabilities under extended 3D conditioning. Together, these data support a qualitative coupling between the strength of ABC transporter programs and 3D-induced functional tolerance, while also highlighting mechanism-specific exceptions that likely reflect pathway rewiring rather than global barrier effects or efflux-driven resistance.

### Growth Kinetics and Gradient Effects Motivate GR-Normalization and Metadata Standards

3DCC size and age influenced both proliferation and apparent pharmacology responses. In our experiments, PANC-1 3DCC continued to expand through day 15 with a peak interval growth rate around day 12 before tapering (Supplementary Figure S3). These evolving baseline growth dynamics illustrate how apparent drug potency and efficacy can be inflated or diminished by changes in proliferation rate as spheroids mature. Because drug-response metrics that do not account for growth kinetics are sensitive to such effects, growth rate–normalized (GR) analyses are essential to distinguish true pharmacologic shifts from proliferation-driven confounding. This normalization is particularly important when comparing responses across culture formats or across timepoints within 3D systems, where growth trajectories are non-linear and state dependent.

Prior literature and our Supplementary Table S7 support a consistent pattern whereby increasing size and age of 3D cellular structures steepen diffusion and oxygen gradients, thereby altering cell-cycle distribution, metabolism, and drug accessibility. Consistent with the emergence of diffusion and oxygen gradients in extended 3D cultures, spheroids exhibited compact architecture and histomorphological features that qualitatively resemble *in vivo* PDAC tissue (Supplementary Figures S4–S6). This sensitivity of pharmacologic measurements to growth state and microenvironmental gradients underscores the importance of explicitly documenting experimental context alongside drug-response data.

At minimum, we recommend reporting the following metadata with 3D drug-response studies (i) pre-culture duration in 3D prior to dosing; (ii) matrix identity/composition (and stiffness when available); (iii) seeding density and aggregate size at the time of dosing ; (iv) media exchange schedule and oxygenation/hypoxia conditions; (v) plate layout/randomization strategy; and (vi) the primary GR metric used (e.g., GR_AOC) together with confidence intervals and replicate structure. Adoption of such reporting standards will facilitate fair cross-study comparison, improve reproducibility, and enhance interpretability across 3D screening platforms.

## Discussion

Extended residence in ECM-containing 3D culture reveals drug-response phenotypes in PDAC models that closely approximate clinical pharmacology, exposing a major limitation of conventional preclinical assays. Conventional systems frequently overestimate drug efficacy for agents that provide minimal benefit to patients with PDAC [5,15,19,49,51–54]. Both 5-fluorouracil (5-FU) and gemcitabine exert pronounced antiproliferative effects in vitro, even at sub-physiological concentrations, yet neither meaningfully improves outcomes for patients with PDAC [55–57]. Such artificial sensitivity, observed in 2D monolayers and patient-derived organoids (PDOs), reflects model-dependent artifacts rather than true therapeutic potential. In contrast, the extended 3D tumoroid systems studied here, MiaPaCa-2, PANC-1, and CFPAC-1, recapitulate resistance levels that more closely align with clinical pharmacology [58,59], consistent with prior studies using hyaluronan-based or ECM-functionalized scaffolds and tumoroid models, that incorporate, hypoxia gradients, multicellular organization, and stromal-like constraints [60–63].

We propose that prolonged residence in ECM-containing 3D culture establishes a “mature tumoroid” state, an experimentally tractable intermediate between short-term spheroids and PDOs. Allowing tumor cells to adapt over time within a defined 3D microenvironment promotes structural compaction, diffusion and oxygen gradients, and metabolic and transcriptional remodeling, thereby activating endogenous resistance programs (including ABC proteins) that mirror those observed in vivo [60,64–70]. In this framework, “Mature Tumoroids” provide a standardized platform to dissect how time-dependent microenvironmental conditioning shapes drug response, bridging the gap between oversensitive 2D assays and heterogeneous patient-derived systems [61,62]. Here, “mature” refers to time-dependent microenvironmental conditioning within a defined 3D ECM context, rather than cellular differentiation or lineage maturation, and does not imply terminal states or loss of plasticity.

Because both 5-FU and gemcitabine have well-characterized clinical pharmacokinetics, they offer a benchmark for evaluating translational fidelity [20–24,71–74]. To place these measurements in clinical context, we compared representative IC₅₀ values across model systems with reported plasma exposure levels for 5-FU (Cmax ≈ 426 µM). Extended 3D tumoroids exhibited IC₅₀ values approaching clinically relevant exposure ranges, whereas conventional 2D cultures substantially underestimated resistance (Figure 8). In our extended 3D cultures, GR₅₀ values closely approximated these physiological exposures. falling within ∼4-fold for 5-FU and ∼0.5-fold for gemcitabine, whereas 2D and short-term 3D systems markedly underestimated resistance. Across nearly all comparisons, extended 3D conditioning produced drug-response predictions that were substantially closer to clinical exposure limits, indicating that the resistance observed here reflects biologically relevant therapeutic ceilings rather than experimental artifacts. Our results establish time-in-3D as a major determinant of PDAC pharmacologic response. When MiaPaCa-2, PANC-1, and CFPAC-1 were pre-cultured in an ECM hydrogel for ≥ 10 days, GR-normalized sensitivity to multiple standards of care declined by one to two orders of magnitude, following a reproducible susceptibility hierarchy (MiaPaCa-2 > PANC-1 > CFPAC-1). Importantly, this tolerance was not uniform: CFPAC-1 displayed increased sensitivity to SN38 and trametinib under extended 3D conditioning, demonstrating that microenvironmental adaptation can invert, rather than simply attenuate, drug responses in a mechanism-dependent manner. Transcriptomic profiling revealed coordinated induction of ABC drug transporters during prolonged 3D residence, with ABCC family members—particularly ABCC3—showing strong association with the functional tolerance observed in extended 3D cultures [31,33,34,75]. Notably, this transporter induction occurred in the absence of prior drug exposure, raising the question of whether endogenous microenvironmental pressures rather than xenobiotic selection drive this phenotype.

**Figure 8.**
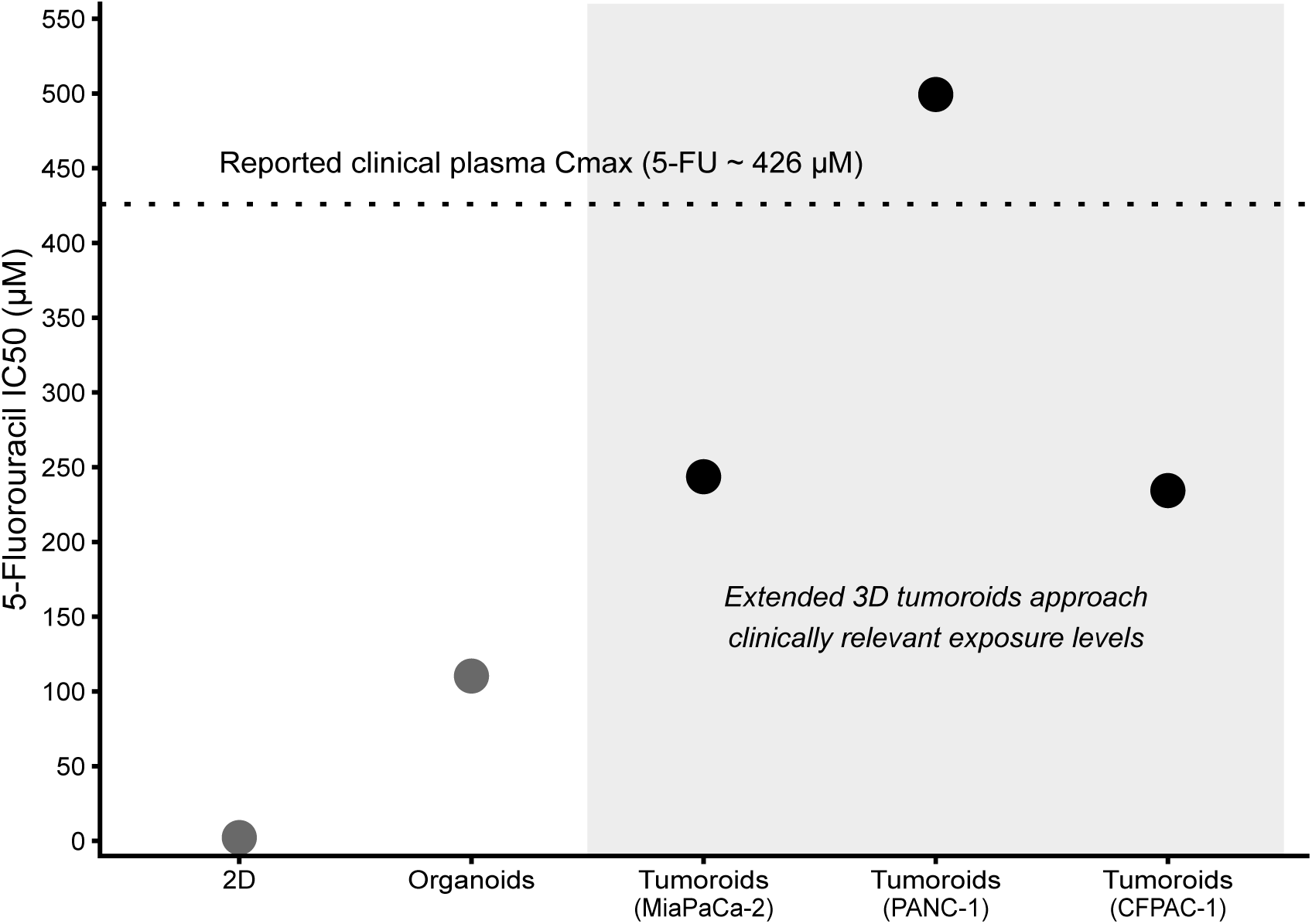
Descriptive comparison of 5-fluorouracil sensitivity across PDAC model systems relative to clinical exposure. IC₅₀ values for 5-fluorouracil measured in 2D cultures, organoids, and extended 3D tumoroid models are displayed relative to reported peak plasma concentrations (Cmax) observed in patients receiving systemic 5-FU therapy. The dashed line indicates the reported clinical plasma Cmax (∼426 µM). This plot is intended as a visual comparison of representative IC₅₀ values across model systems rather than a statistical analysis. Extended 3D tumoroids exhibit IC₅₀ values approaching clinically relevant exposure levels, whereas conventional 2D cultures substantially underestimate drug resistance.

ABCC3 is frequently upregulated in PDAC and has been linked to reduced sensitivity across multiple drug classes, consistent with the rightward GR shifts observed here. Mechanistically, ABCC3 mediates efflux of conjugated metabolites and couples to detoxification pathways, thereby reducing effective intracellular drug exposure and blunting cytotoxic responses [31,33,34,75–80]. In parallel, ABCA8 represents a distinct, signaling-competent resistance axis: by exporting taurocholic acid and activating S1PR2–ERK signaling, ABCA8 can attenuate gemcitabine response through mechanisms extending beyond classical drug efflux [34]. Although our analyses are primarily transcript-based, the concordance between the functional resistance map (ΔGR_AOC) and transporter induction motivates targeted genetic or acute pharmacologic perturbations to test necessity and sufficiency within the pre-conditioned 3D context. Interestingly, the emergence of ABC transporter expression in extended 3D cultures occurred in the absence of prior drug exposure, suggesting that mechanisms beyond classical xenobiotic selection may contribute to this phenotype. One possible explanation involves metabolic adaptation to the constrained microenvironment of dense three-dimensional tumor structures. Restricted diffusion, hypoxia, and altered nutrient gradients are known to reprogram cellular metabolism and may promote accumulation of potentially toxic endogenous metabolites [81]. In this context, ABC transporters may function not only as drug efflux systems but also as exporters of endogenous metabolic byproducts, enabling tumor cells to maintain metabolic homeostasis within confined architectures. Consistent with this interpretation, when mature 3D cultures were dissociated and replated, a substantial portion of the resistance phenotype diminished (data not shown), suggesting that transporter induction may represent a context-dependent and partially reversible adaptation to prolonged 3D residence.

The sensitivity inversions observed in CFPAC-1 are particularly informative, indicating that extended microenvironmental conditioning does not simply shift tumor cells along a one-dimensional resistance axis. Instead, prolonged residence in a defined 3D microenvironment can rewire pathway dependencies in a context- and lineage-dependent manner, exposing vulnerabilities that are not detected in short-term 2D or 3D assays [62,82]. These inversions highlight the risk of overlooking actionable liabilities when pre-culture duration is ignored and provide tractable entry points for mechanistic dissection of ERK signaling, DNA-damage response, or topoisomerase engagement under mature 3D conditions.

Methodologically, growth-rate (GR)–based analysis was essential to disentangle true pharmacologic shifts from proliferation confounding across formats and timepoints [41,45]. GR approaches enable fair comparisons but require careful implementation, including accurate baseline growth estimation, robust curve fitting, and transparent reporting of effect sizes. We recommend reporting full dose–response curves with confidence intervals, a pre-specified primary GR metric (e.g., GR_AOC) for activity calls, effect sizes with uncertainty estimates (ΔGR_AOC or GR₅₀ shifts), and appropriate multiplicity control across multi-drug panels. Where feasible, orthogonal readouts, such as live-cell imaging, label-free confluence tracking, or apoptosis markers, should be used to corroborate fluorescence-based viability measurements in 3D matrices.

To enhance reproducibility and cross-study comparability, we recommend routine disclosure of key 3D metadata, including pre-culture duration prior to dosing, matrix identity and mechanical properties (e.g., stiffness), seeding density and aggregate size at the time of treatment, media-exchange schedules, oxygenation or hypoxia conditions, and plate layout and randomization [39,44,83,84]. Because time-in-3D modulates diffusion distances, oxygen gradients, and cell-cycle state, these parameters should be logged alongside dose and exposure time to enable fair benchmarking and meta-analysis.

From a translational perspective, extended 3D conditioning more closely models tumor residence within desmoplastic stromal niches, where sustained ECM exposure, hypoxia, and metabolic stress shape therapeutic response [85–88]. If the tolerance observed here reflects a durable, inducible program, intervention timing becomes critical: agents targeting efflux or hypoxia-associated pathways may be most effective after defined pre-conditioning intervals, whereas fast-acting cytotoxic may benefit from strategies that disrupt or prevent tolerance induction. In parallel, the broad dysregulation of ABC transporters in PDAC—including ABCB4, ABCB11, ABCC1, ABCC3, ABCC5, ABCC10, ABCG2, and lipid-handling members ABCA1, ABCA7, and ABCG1—supports the development of transporter-linked biomarkers to guide patient stratification and sequence-dependent combination strategies [30,88–95]. Collectively, these data establish time-in-3D as a first-order determinant of PDAC pharmacology. Extended 3D residence broadly induces tolerance in MiaPaCa-2, and PANC-1 yet simultaneously reveals mechanism-specific vulnerabilities in CFPAC-1, including enhanced sensitivity to SN38 and trametinib. The coordinated up-regulation of ABC transporters after prolonged 3D conditioning provides a plausible mechanistic substrate for reduced intracellular drug exposure and blunted cytotoxicity, while the inversion cases argue for focused pathway-level follow-up, including MEK/ERK signaling and DNA-damage or topoisomerase engagement. Practically, these findings support: (i) incorporating both short- and extended-3D pre-conditioning arms in preclinical and PDO studies; (ii) routine reporting of pre-culture duration alongside GR-normalized effect sizes; and (iii) exploration of sequencing strategies that interleave efflux- or hypoxia-modulating agents with cytotoxic to counter inducible tolerance.

This study has limitations, including reliance on established cell lines, transcript-level associations without direct causality testing, and limited spatial pharmacokinetic measurements. Addressing these limitations through targeted transporter perturbations, intra-spheroid drug-penetration assays, expanded ECM chemistries, and integration of patient-derived organoids with single-cell or spatial readouts will strengthen generalizability and translational anchoring [83,84].

In summary, time-dependent microenvironmental conditioning emerges as a dominant but under-controlled determinant of PDAC drug response. Extended 3D residence broadly induces tolerance while unmasking mechanism-specific vulnerabilities in defined contexts. Recognizing and standardizing this variable, alongside culture format and dose, will refine preclinical benchmarking and improve alignment between in vitro pharmacology and clinical behavior in PDAC.

## Materials and Methods

### Cell culture

CFPAC-1, MiaPaCa-2, and PANC-1 pancreatic ductal adenocarcinoma cell lines were obtained from the European Collection of Authenticated Cell Cultures (ECACC, UK). All lines were maintained in Advanced DMEM/F-12 medium (Gibco, Europe) supplemented with 5% fetal bovine serum (EURx, Poland) and 1× GlutaMAX (Gibco, Europe). Cells were cultured at 37 °C in a humidified atmosphere containing 5% CO₂.

For both 2D and 3D assays, cells were harvested using TrypLE Express (Santa Cruz Biotechnology, Germany) and resuspended in complete culture medium. Cells were seeded at a density of 500 cells per well in a total volume of 60 µL into 384-well plates.

For 2D culture, cells were plated onto tissue-culture–treated, flat-bottom 384-well plates (Greiner Bio-One, product no. 781182) and allowed to attach overnight prior to initiation of drug treatment.

For 3D culture, cells were seeded onto hydrogel-containing 384-well plates (LifeGel, product RRLG384-A2G; Real Research, Poland). Plates were pre-equilibrated with culture medium, after which excess medium was aspirated to leave a thin liquid layer above the hydrogel surface before cell seeding. Cells were allowed to attach and grow under 3D conditions for 1–12 days prior to drug treatment. All 3D cultures were maintained at 37 °C and 5% CO₂ throughout the pre-culture period.

### Drug treatment

Stock solutions of obatoclax (Selleck GmbH, Germany), 5-fluorouracil (5-FU), and 7-ethyl-10-hydroxycamptothecin (SN38) were prepared at 10 mM in dimethyl sulfoxide (DMSO). Doxorubicin hydrochloride (Sigma, Merck Life Science, Poznań, Poland) was prepared as a 5 mM stock in DMSO. Gemcitabine hydrochloride (TCI, Japan) was prepared as a 10 mM stock in water, while oxaliplatin and carboplatin (MedChemExpress, Monmouth Junction, NJ, USA) were prepared as 5 mM aqueous stock solutions. All remaining compounds were purchased from MedChemExpress as ready-to-use 10 mM stock solutions in DMSO.

All drug stocks were stored at −80 °C until use. On the day of treatment, 3-fold serial dilutions were prepared in complete culture medium. For DMSO-based compounds, DMSO concentration was kept constant across all dilutions, resulting in a final well concentration of 0.1% DMSO. Gemcitabine, oxaliplatin, and carboplatin dilutions contained no DMSO.

Prior to drug addition, culture medium was aspirated from both 2D and 3D wells, leaving approximately 25 µL of residual medium or medium–hydrogel volume. Drug-containing or control medium (40 µL) was then added to each well. Reported drug concentrations reflect the final concentration following diffusion into a total well volume of approximately 65 µL (medium plus hydrogel). All drug treatments were carried out for 72 hours, after which cell viability assays were performed.

For 2D experiments, drug treatment was initiated one day after cell seeding. For comparative 3D experiments, drug treatment was initiated after defined pre-culture intervals corresponding to early (day 4–7) and extended (‘mature’) conditions (day 10–12), depending on the experimental design. For clarity, results are reported using representative timepoints (day 7 and day 11), which capture early and mature 3D states, respectively. For experiments involving drug addition at day 12, medium exchanges were performed on days 8 and 10 prior to treatment to maintain nutrient availability during prolonged 3D culture. Compounds, concentrations, and reported mechanisms of action are summarized in Supplementary Table S8.

### Cell viability measurements

Cell viability was quantified using the CellTiter-Blue® assay (Promega, Madison, WI, USA), which measures metabolic reduction of resazurin to the fluorescent product resorufin. Assay conditions were optimized to maintain linearity of the fluorescence signal with respect to both cell number and incubation time across all growth schedules and culture durations.

To account for differences in metabolic activity associated with extended culture and spheroid maturation, assay parameters were adjusted by modulating either the volume of CellTiter-Blue® reagent added per well and/or the incubation time. For assays performed up to 7 days after initial seeding, 7.5 µL of CellTiter-Blue® reagent was added per well. For assays performed after ≥8 days of culture, 15 µL of reagent was used to ensure sufficient signal intensity without saturation.

Following reagent addition, plates were incubated at 37 °C in a humidified atmosphere containing 5% CO₂ for 0.5–4 hours, depending on the specific growth condition. Fluorescence was measured using a VICTOR Nivo multimode plate reader (Revvity, Waltham, MA, USA) with excitation and emission wavelengths of 570 nm and 615 nm, respectively.

To enable calculation of growth rates for GR-based analysis, baseline (pre-treatment) viability measurements were performed on each assay plate immediately prior to drug addition using the same assay conditions. These measurements provided the reference values required for normalization of drug-response data to proliferation rate across culture formats and time points.

### Immunohistochemical staining of FFPE 3D cell culture samples

Formalin-fixed, paraffin-embedded (FFPE) 3D cell culture samples were sectioned at 4–5 µm thickness and mounted onto glass microscope slides. Slides were warmed at 46 °C for 30 min to promote section adhesion. Sections were then deparaffinized in xylene (2 × 15 min) and rehydrated through a graded ethanol series (100%, 96%, 80%, 70%, and 50%; 5 min each) to distilled water.

For histological staining, sections were immersed in Mayer’s hematoxylin for 5 min, rinsed thoroughly in distilled water, and washed under running tap water for 10 min to allow complete bluing. Sections were subsequently counterstained with eosin Y (0.5–1%) for 1 min, followed by dehydration through ascending ethanol concentrations (70%, 80%, 96%, and 100%; 2 × 5 min each) and clearing in xylene (2 × 5 min).

Slides were mounted using Consul-Mount resin-based mounting medium and sealed with glass coverslips. After air drying, sections were imaged by brightfield microscopy using identical illumination and exposure settings across samples to ensure comparability.

### Proteomic sample preparation

Cell pellets from 2D cultures were lysed directly in lysis buffer (8 M urea, 30% acetonitrile, 100 mM NH₄HCO₃); pellets from 3D cultures were first recovered by collagenase/hyaluronidase digestion followed by mechanical homogenization. Lysates were sonicated (20 min), vortexed (30 min), and clarified by centrifugation at 17,000 × g for 30 min. Protein concentration was determined with the Qubit Protein Broad Range Assay (Invitrogen), and 60 µg of total protein per sample was carried forward.

### In-solution digestion

Proteins were reduced with 20 mM DTT (30 min, 37 °C), alkylated with 60 mM IAA (30 min, RT, dark), and diluted to 1 M urea with 100 mM NH₄HCO₃. Digestion was performed overnight at 37 °C with 2 µg sequencing-grade trypsin (Promega) and quenched with 5% TFA. Peptides were desalted on C18 Micro SpinColumns (Harvard Apparatus), eluted with 0.1% TFA/80% acetonitrile, and dried by SpeedVac at 30 °C.

### LC–MS/MS acquisition

Dried peptides were reconstituted in 30 µL of 0.08% TFA/2.5% acetonitrile, quantified by NanoDrop at 220 nm, and spiked with iRT standards (Biognosys). Samples were analyzed on an UltiMate 3000 RSLCnano system coupled to an Orbitrap Exploris 480 (Thermo Scientific). Peptides were trapped on a C18 PepMap column (0.3 × 5 mm) and separated on an analytical C18 PepMap RSLC column (75 µm × 250 mm, 2 µm) using a 90-min linear gradient from 2.5% to 40% acetonitrile (0.1% formic acid). For each sample, one DDA run and two DIA technical replicates were acquired.

DDA: full MS at 120,000 resolution (m/z 350–1200; AGC 300%, auto injection time); MS/MS at 30,000 resolution (HCD 30%, 2 m/z isolation, charge states 2–6, intensity threshold 5,000; dynamic exclusion 20 s, ±10 ppm).

DIA: full MS at 60,000 resolution (m/z 350–1450; AGC 300%, 100 ms injection time); 62 × 12 m/z isolation windows (1 m/z overlap, 350–1100) with fragment acquisition at 30,000 resolution (HCD 30%, AGC 1000%). All spectra stored in profile mode.

### Spectral library generation and database search

Human and murine FASTA databases (UniProt Swiss-Prot + TrEMBL, accessed 15 November 2023) were merged after removal of shared sequences; contaminant and decoy sequences were appended. Raw files were converted to mzML with MSConvert v1.5.2 (peak picking enabled). Three complementary spectral libraries were generated and combined: (i) a DDA-based library and (ii) a pseudo-DDA library from DIA-Umpire signal extraction, both using FragPipe v20.0; and (iii) an in-silico library from DIA-NN v1.8.1. Retention times were aligned with a custom Python script before merging in DIA-NN. The consensus library was filtered to human-specific precursor peptides.

FragPipe parameters: precursor ±15 ppm, fragment 15 ppm; strict trypsin; ≤2 missed cleavages; fixed carbamidomethylation (C); variable oxidation (M), protein N-terminal acetylation, and N-terminal Met excision (≤3 variable modifications, ≤5,000 combinations); charge 1–4; peptide length 7–50 aa; PSM validation by Percolator.

DIA-NN parameters: trypsin/P; 1 missed cleavage; variable modifications as above (≤1); peptide length 7–30 aa; precursor charge 1–4, m/z 300–1800; fragment m/z 200–1800; precursor FDR 1%; mass accuracy 17 ppm, MS1 accuracy 16 ppm; scan window 11; isotopologues, MBR, and no shared spectra enabled; species-specific protein inference; double-pass neural network; robust LC quantification; RT-dependent cross-run normalization.

### Data analysis

Peptide-level quantification matrices were extracted from DIA-NN output using the diann R package; precursor intensities were summarized to peptide level with MaxLFQ and subsequently aggregated to gene level using the QFeatures R package. Technical DIA replicates were merged, low-quality features filtered, and the resulting gene-level matrix was quantile-normalized to correct for systematic technical variation.

### Drug response quantification and GR-based analysis

Raw fluorescence intensity values from cell viability assays were first corrected by subtraction of no-cell background signal. Vehicle-treated (no-drug) control wells were normalized to 100% viability, and percent viability was calculated for each drug concentration.

Drug dose–response curves were fitted using the Growth Rate (GR) analysis framework implemented at http://www.grcalculator.org [45]. This approach yields conventional IC₅₀ values as well as GR-corrected metrics, including GR₅₀ (the concentration causing 50% growth-rate inhibition) and GR_AOC (growth-rate area over the curve), enabling comparisons across conditions with differing proliferation rates.

For each drug–condition pair, mean IC₅₀, GR₅₀, and GR_AOC values were calculated from at least three independent curve fits corresponding to technical replicates. Each dose–response curve consisted of 18 data points, derived from nine drug concentrations tested in duplicate.

Statistical comparisons of GR_AOC values between conditions were performed using a two-sample *t*-test assuming equal variance, implemented in Microsoft Excel. Statistical significance thresholds are indicated in the corresponding figures and tables.

