## Supplemental FIles for "Mature tumoroids recapitulate clinically relevant drug response through extended 3D culture in PDAC"

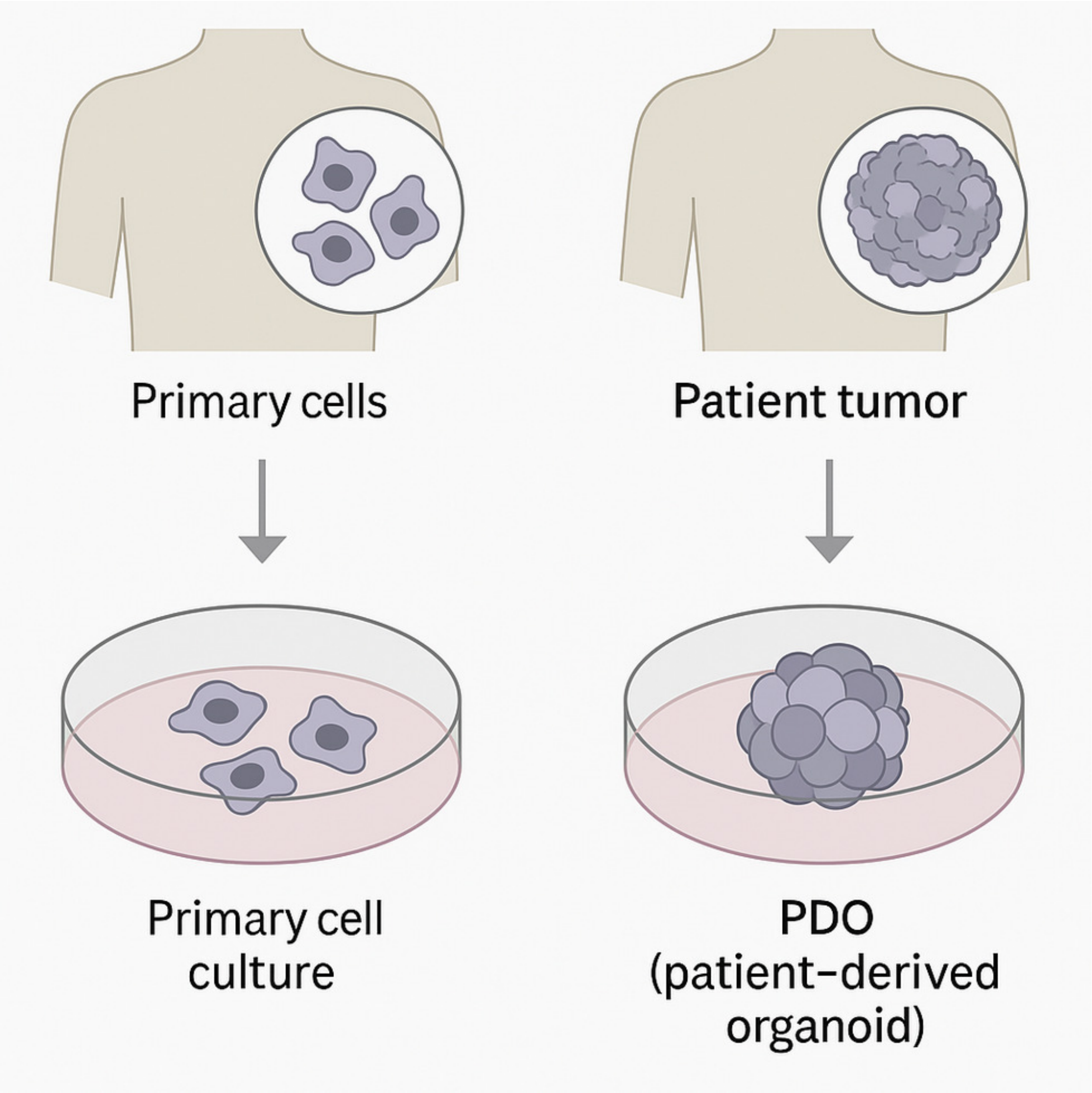


**Supplementary Figure S1 | Comparison between primary cell cultures and patient-derived organoids (PDOs).**

Primary cells are isolated from tissue and cultured as dissociated cells in a 2D monolayer, which does not maintain native tissue architecture over prolonged culture. In contrast, patient tumor samples retain cellular heterogeneity and are embedded in a 3D matrix (e.g., Matrigel) to generate patient-derived organoids (PDOs) that self-organize into structures recapitulating key aspects of the original tumor’s architecture, mutational landscape, and functional characteristics.

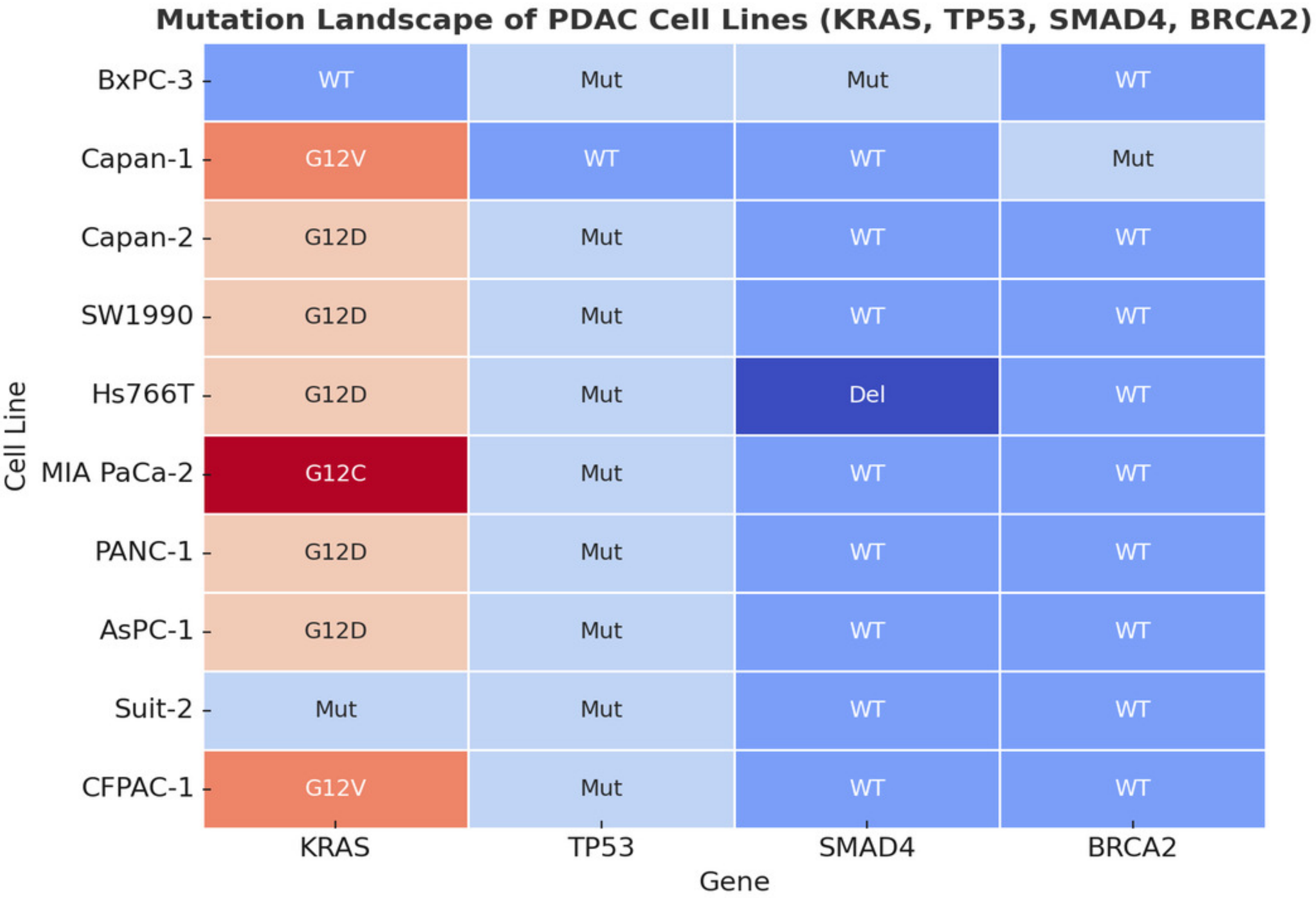


**Supplementary Figure S2 | Mutation landscape of commonly used PDAC cell lines.**

Heatmap summarizing the mutational status of key pancreatic ductal adenocarcinoma (PDAC) driver genes—**KRAS, TP53, SMAD4, and BRCA2**—across representative PDAC cell lines. Colors indicate wild-type (WT), mutant (Mut), or specific alterations where indicated (e.g., KRAS G12D, G12V, G12C; SMAD4 deletion). This overview provides genomic context for the cell lines analyzed in this study but is not intended to imply direct causal relationships with drug-response phenotypes.

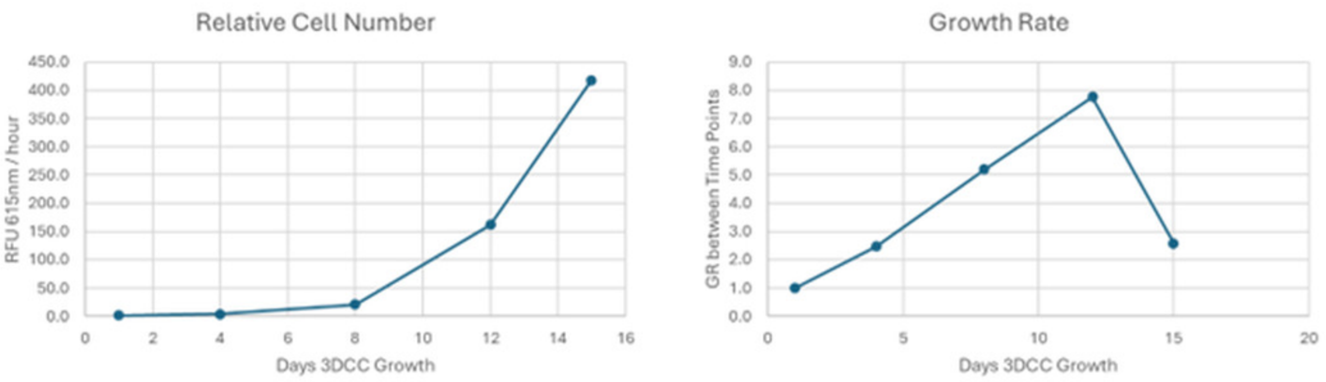


### Supplementary Figure S3 | PANC-1 proliferates during extended 3D pre-culture.

**Left:** Longitudinal increase in relative cell number (metabolic RFU per well) during culture in an ECM-containing 3D hydrogel, indicating continued expansion through day 15.

**Right:** Interval growth rate (GR) calculated between successive time points. GR increases during early–mid 3D culture, peaks around day 12, and modestly tapers by day 15 as spheroids enlarge. These growth dynamics define the baseline proliferation behavior used for GR-normalized drug-response analyses.

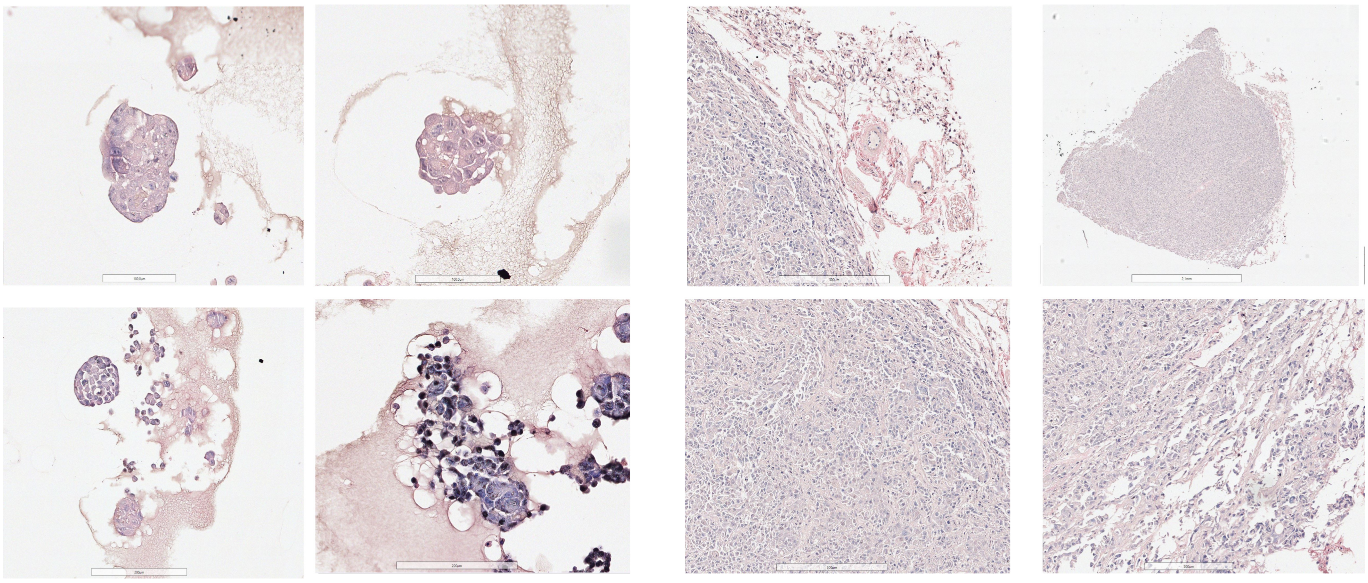

**Supplementary Figure S4 | Morphological comparison of PANC-1 spheroids cultured in 3D LifeGel and in vivo tumor tissue.**

Representative histological images of **PANC-1 pancreatic cancer cells** grown in a **defined 3D LifeGel hydrogel** (left) compared with **PANC-1–derived tumors in mouse tissue** (right). In 3D LifeGel, PANC-1 cells form **compact, multilayered spheroid structures with limited organization**, reflecting a poorly differentiated, mesenchymal-like phenotype. In vivo tumor sections display **dense, solid tumor architecture with stromal infiltration and minimal glandular features**, closely resembling the morphology observed in extended 3D cultures. These qualitative similarities support the notion that prolonged culture in a defined 3D ECM promotes **architecture-dependent tumor states** that better approximate in vivo growth patterns compared with 2D monolayers. Scale bars as indicated.

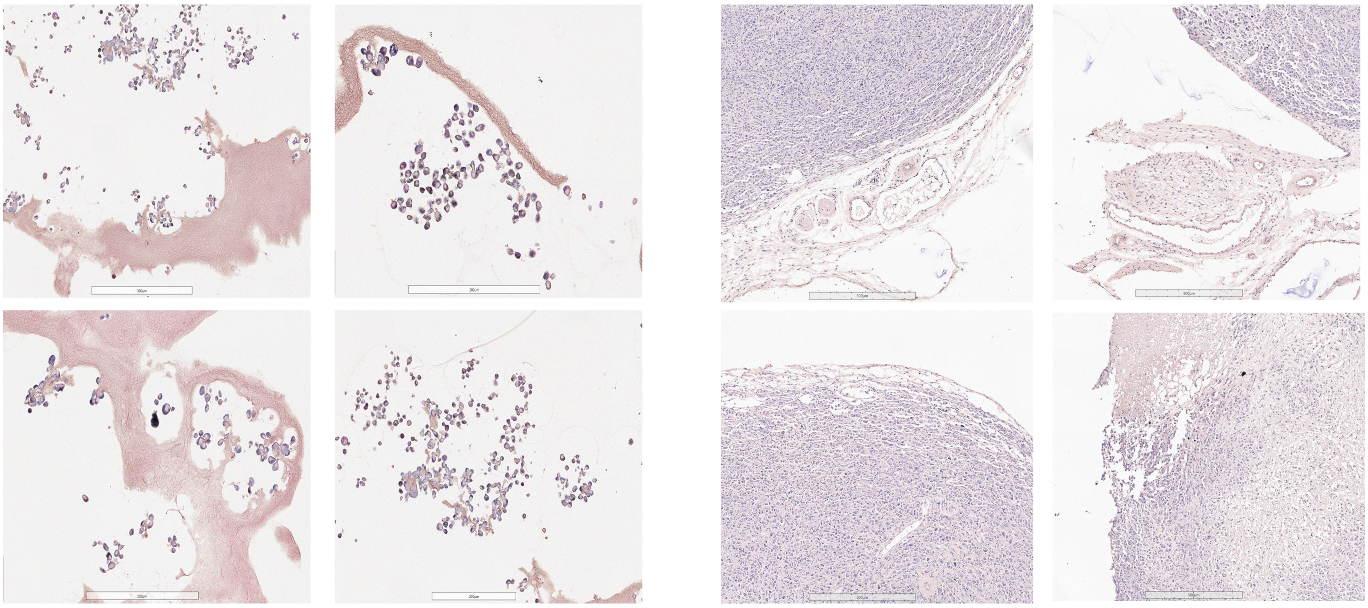

**Supplementary Figure S5 | Morphological comparison of MiaPaCa-2 spheroids cultured in 3D LifeGel and in vivo tumor tissue.**

Representative histological images of **MiaPaCa-2 pancreatic cancer cells** grown in a **defined 3D LifeGel hydrogel** (left) compared with **MiaPaCa-2–derived tumors in mouse tissue** (right). In 3D LifeGel, MiaPaCa-2 cells predominantly form **compact, solid aggregates with limited lumen formation**, reflecting a poorly differentiated architecture. In vivo tumor sections display **dense, solid tumor organization with minimal glandular features**, closely resembling the morphology observed in extended 3D cultures. This qualitative comparison indicates that prolonged 3D ECM culture promotes **architecture-dependent phenotypes** in MiaPaCa-2 cells that resemble in vivo tumor structure. Scale bars as indicated.

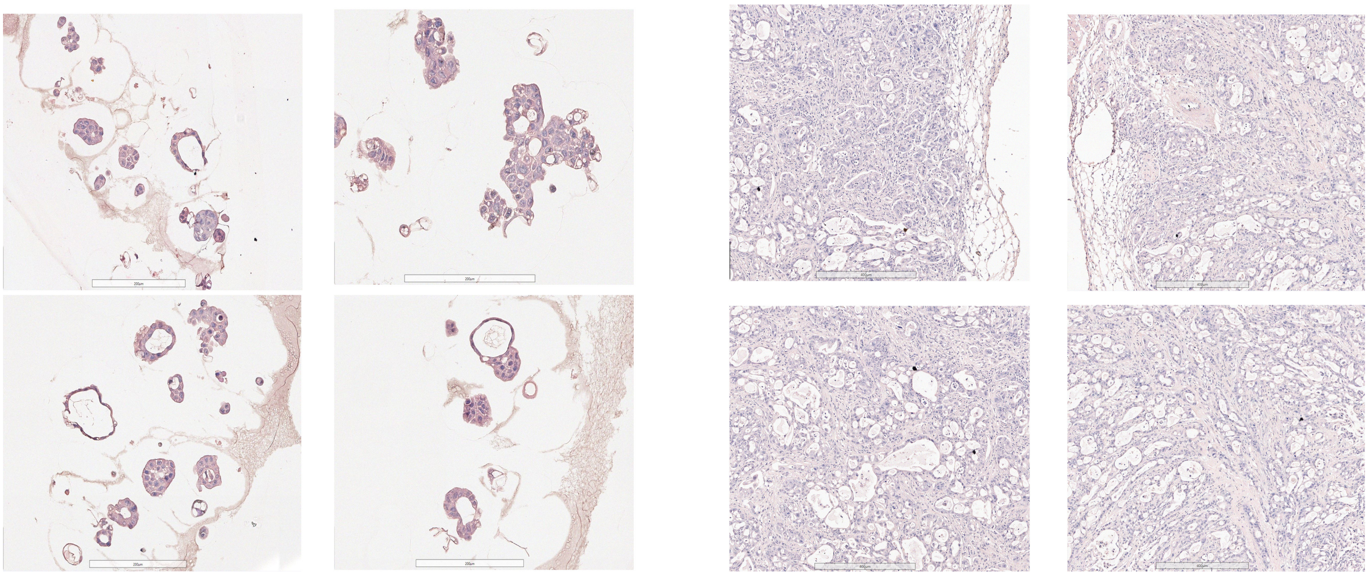

**Supplementary Figure S6 | Morphological comparison of CFPAC-1 spheroids cultured in 3D LifeGel and in vivo tumor tissue.**

Representative histological images of **CFPAC-1 pancreatic cancer cells** grown in a **defined 3D LifeGel hydrogel** (left) compared with **CFPAC-1–derived tumors in mouse tissue** (right). 3D LifeGel cultures form **cyst-like epithelial structures with luminal organization**, reminiscent of glandular features observed in vivo. Tumor sections show **dense epithelial architecture with stromal interfaces**, reflecting the native tumor microenvironment.This comparison illustrates that CFPAC-1 cells adopt **architecture-dependent morphological features in 3D culture** that qualitatively resemble in vivo tumor organization, supporting the physiological relevance of extended 3D ECM models. Scale bars as indicated.

### Supplementary Table S1 | Comparison between primary cell cultures and patient-derived tumor samples used for organoid generation (PDOs).

| Aspect | Primary Cells | Patient Tumor Samples (for PDOs) |
|---|---|---|
| **Source** | Isolated from **healthy or diseased tissue** (e.g., normal pancreas, liver, skin, or even a tumor), but **cultured as a single-cell population** in the lab. | Directly taken from a **patient's tumor biopsy or surgical specimen**, containing **multiple cell types** (cancer cells + stromal + sometimes immune cells) and preserving **tumor architecture**. |
| **Processing** | Typically **dissociated into individual cells** and cultured under specific conditions to maintain their phenotype for a short period. They often lose tissue architecture quickly. | Processed to **retain 3D structure and cellular heterogeneity:** small fragments or dissociated cells are embedded in a supportive matrix (like Matrigel) to self-organize into organoids that mimic the tumor's original structure. |
| **Lifespan in culture** | Usually **short-lived** — they undergo senescence after a few passages because they are not immortalized. | Can often be **propagated long-term** as organoids while maintaining key genetic and histological features of the original tumor. |
| **Genetic background** | May be **normal (non-cancerous)** or partially altered, depending on the donor and isolation. | Largely reflects the patient's cancer genotype, including driver mutations, chromosomal rearrangements, and intra-tumoral heterogeneity. |
| **Complexity** | Homogeneous — mainly one cell type. | Heterogeneous: multiple interacting cell types (epithelial, stromal, etc.) in a self-organized 3D structure. |

Primary cell cultures and patient-derived organoids differ in their source, processing, lifespan in culture, genetic background, and biological complexity. Primary cells are typically cultured as dissociated cells under 2D conditions and provide simplified, short-term models, whereas patient tumor samples embedded in 3D matrices can self-organize into organoids that retain key architectural, genetic, and phenotypic features of the original tumor. This comparison highlights the complementary roles of these systems in cancer modeling and translational research.

### Supplementary Table S2 | Comparison of engineered 3D models and patient-derived organoids (PDOs).

| Feature | Engineered 3D models (defined matrices, synthetic hydrogels) | Patient-derived organoids (PDOs, Matrigel-based) |
|---|---|---|
| **Origin** | Established cell lines or primary cells embedded in a defined biomaterial (e.g., PEG, alginate, collagen, tunable ECM mixtures). | Directly derived from patient tumour samples maintaining endogenous architecture and mutations. |
| **Matrix definition** | Chemically and mechanically defined; batch-to-batch reproducible tunable stiffness and ligand presentation. | Typically Matrigel or basement-membrane extract; complex and partially undefined composition with batch-to-batch variability. |
| **Control of microenvironmental cues** | High stiffness, ECM composition, oxygen and nutrient diffusion can be independently modulated. | Limited independent control over microenvironmental parameters, which are largely dictated by matrix composition and culture conditions. |
| **Reproducibility** | High experimental reproducibility and scalability for screening. | Moderate; inter- and intra-patient variability is intrinsic and biologically meaningful but limits standardization. |
| **Physiological relevance** | Captures specific physical and biochemical cues of the tumour microenvironment; allows mechanistic dissection. | Recapitulates genomic, transcriptomic and phenotypic heterogeneity of patient tumours. |
| **Drug-response interpretation** | Enables quantitative, mechanism-focused assays under standardized conditions. | Reflects clinical variability but quantitative interpretation can be complicated by matrix effects and biological heterogeneity. |
| **Applications** | Mechanistic studies, microenvironmental perturbation, controlled drug-response modeling, bioengineering. | Precision medicine, inter-patient response prediction, biobanking, personalized drug testing. |
| **Limitations** | Limited genetic heterogeneity; may not fully capture tumour evolution or stromal complexity. | Complex and partially undefined ECM, variable growth rates, and limited independent control over physical cues. |
| **Representative studies** | [11] Gjorevski et al., 2016 (Nature);<br>[18] Hoarau-Villiers et al., 2023 (Commun. Biol.). | [14] Sachs et al., 2018 (Cell).<br>[15] Tiriac et al., 2018 (Cancer Discov.); |

Engineered 3D models based on defined matrices enable systematic manipulation of biophysical and biochemical microenvironmental parameters with high reproducibility, supporting quantitative and mechanistic drug-response studies. In contrast, patient-derived organoids preserve the genetic and phenotypic diversity of human tumours but exhibit intrinsic variability associated with complex and partially undefined matrix environments. Together, these complementary systems provide a robust framework for investigating therapeutic responses and tumour microenvironmental determinants.

5 out of 5 standard of care drugs (all 14 test drugs) remain active after 7 days 3D growth ($p < 0.05$)
2 out of 5 standard of care drugs (11 of 14 total test drugs) remain active after extended 3D growth ($p < 0.05$)

| | Panc-1 2D $IC_{50}$ Day4 | Panc-1 3D $IC_{50}$ Day7 | Panc-1 3D $IC_{50}$ Day11 | Panc-1 2D $GR_{50}$ Day4 | Panc-1 3D $GR_{50}$ Day7 | Panc-1 3D $GR_{50}$ Day11 | Panc-1 2D GR_AOC Day4 | Panc-1 3D GR_AOC Day7 | Panc-1 3D GR_AOC Day11 |
|---|---|---|---|---|---|---|---|---|---|
| 5-FU | 15.9 | 16.9 | 340 | 6.39 | 9.01 | 105 | 1.12 | 0.98 | 0.32 * |
| GEM | NF | 5.98 | > 10 | NF | 2.87 | 30.1 | 0.44 | 0.60 | 0.58 |
| OXA | 1.63 | 8.32 | 13.6 | 0.592 | 2.34 | 4.72 | 1.32 | 0.68 | -0.15 ** |
| PAC | 1.98E-06 | 0.000325 | > 10 | 1.57E-07 | 0.000117 | 0.186 | 5.16 | 4.36 | 1.50 * |
| SN38 | 0.212 | 0.188 | 2.06 | 0.0921 | 0.0965 | 0.367 | 2.26 | 3.16 | 1.81 |
| BOR | 0.00511 | 0.00490 | 0.0142 | 0.00456 | 0.00329 | 0.00891 | 4.16 | 4.57 | 5.35 |
| CAR | NF | 2.96 | 11.9 | NF | 1.48 | 9.44 | 0.03 | 0.33 | -0.15 |
| DIS | 0.253 | 5.42 | 13.0 | 0.192 | 4.89 | 9.24 | 1.20 | 1.27 | 0.13 |
| DOX | 1.60 | 0.808 | 43.8 | 0.601 | 1.68 | 7.07 | 1.32 | 0.97 | 0.59 |
| HOM | 0.0297 | 0.0316 | 0.0410 | 0.0218 | 0.0192 | 0.0259 | 4.00 | 4.65 | 4.23 |
| KPT | 0.878 | 1.07 | > 10 | 0.459 | 0.442 | > 10 | 1.15 | 1.41 | 0.46 |
| OBA | 0.483 | 0.626 | 2.07 | 0.324 | 0.406 | 0.750 | 1.67 | 1.97 | 1.44 |
| TRA | NF | NF | 25.3 | NF | 0.000595 | 1.09 | 1.90 | 3.05 * | 1.30 |
| YM155 | 0.000688 | 0.00258 | 0.0169 | 0.000597 | 0.00221 | 0.0115 | 5.94 | 6.00 | 5.54 |

$IC_{50}$ / $GR_{50}$ values in uM (determined from independent triplicate curve fits, 18 data points per curve)
AOC = Area Over Curve
Day = Age of culture after 72h drug treatment
reduction or gain in drug sensitivity ($IC_{50}$ or $GR_{50}$ 2D vs 3D value changes > 5x, AOC value changes > 1.5x)

* $p < 0.05$, ** $p < 0.01$

### Supplementary Table S3 | Summary of 2D vs 3D drug-sensitivity metrics for PANC-1.

PANC-1 cells were cultured in 2D (day 4) or in an ECM-containing 3D hydrogel (day 7 or day 11) and treated for 72 h with the indicated agents. Columns report $IC_{50}$ (µM), $GR_{50}$ (µM), and GR_AOC (area over the GR curve; unitless), derived from independent triplicate dose–response fits (18 data points per curve). Values listed as ">10" or ">100" exceeded the highest concentration tested. Lower $IC_{50}$/$GR_{50}$ and lower GR_AOC indicate higher sensitivity. Color shading encodes the direction and magnitude of change relative to 2D day 4 (pink, decreased sensitivity in 3D; green, increased sensitivity in 3D; thresholds: ≥5× change for $IC_{50}$/$GR_{50}$ and ≥1.5× for GR_AOC). Asterisks denote significant differences between 3D and 2D conditions (*$P < 0.05$; **$P < 0.01$).

**Overall activity summary:** All five standard-of-care agents (14/14 total drugs tested) retain activity at 3D day 7 ($P < 0.05$), whereas only two of five standard-of-care agents (11/14 total drugs tested) remain active after extended 3D growth (day 11; $P < 0.05$). Consistent with the GR dose–response curves, PANC-1 cells exhibit broad reductions in drug sensitivity with prolonged 3D residence, with only limited instances of maintained or increased sensitivity.

3 out of 5 standard of care drugs (9 of 14 total test drugs) remain active after 7 days 3D growth ($p < 0.05$)
0 out of 5 standard of care drugs (2 of 14 total test drugs) remain active after extended 3D growth ($p < 0.05$)

| | MiaPaca2 2D $IC_{50}$ Day4 | MiaPaca2 3D $IC_{50}$ Day7 | MiaPaca2 3D $IC_{50}$ Day11 | MiaPaca2 2D $GR_{50}$ Day4 | MiaPaca2 3D $GR_{50}$ Day7 | MiaPaca2 3D $GR_{50}$ Day11 | MiaPaca2 2D GR_AOC Day4 | MiaPaca2 3D GR_AOC Day7 | MiaPaca2 3D GR_AOC Day11 |
|---|---|---|---|---|---|---|---|---|---|
| 5-FU | 2.23 | 9.97 | > 100 | 4.13 | 22.1 | > 100 | 1.44 | 0.88 | 0.00 * |
| GEM | 0.00533 | 0.0137 | > 10 | 0.00743 | 0.518 | > 10 | 3.27 | 1.82 | 0.26 ** |
| OXA | 1.26 | 6.23 | > 10 | 1.45 | 14.5 | > 10 | 1.38 | 0.19 ** | 0.28 ** |
| PAC | 5.96E-07 | 0.0313 | > 10 | 5.96E-07 | 0.0167 | > 10 | 3.77 | 1.83 * | -0.53 ** |
| SN38 | 0.00200 | 0.00430 | 1.60 | 0.0282 | 0.0116 | 0.387 | 4.69 | 4.05 | 1.55 ** |
| BOR | 0.00531 | 0.00910 | 0.797 | 0.00737 | 0.00945 | 0.378 | 3.14 | 2.37 | 2.28 |
| CAR | > 10 | > 10 | > 10 | > 10 | 15.3 | > 10 | 1.07 | 0.00 ** | 0.26 ** |
| DIS | > 10 | 6.05 | > 10 | 11.2 | 7.67 | > 10 | 1.76 | 0.44 ** | 0.01 ** |
| DOX | 0.0670 | 0.351 | 7.32 | 0.445 | 0.978 | 5.37 | 2.60 | 2.19 | 0.23 * |
| HOM | 0.00319 | 0.0106 | 0.0929 | 0.00638 | 0.0162 | 0.0495 | 6.35 | 4.98 | 3.89 ** |
| KPT | 0.0912 | 0.314 | > 10 | 0.489 | 0.988 | > 10 | 2.40 | 2.01 | 0.39 ** |
| OBA | 0.0326 | 0.263 | 4.77 | 0.189 | 1.36 | 3.21 | 3.42 | 1.89 ** | 0.83 ** |
| TRA | 5.96E-07 | 5.96E-07 | 0.0467 | 5.96E-07 | 5.96E-07 | 0.00498 | 5.55 | 9.22 ** | 4.87 |
| YM155 | 0.00238 | 0.00737 | 0.959 | 0.00412 | 0.0118 | 0.426 | 4.70 | 3.83 | 2.37 ** |

$IC_{50}$ / $GR_{50}$ values in uM (determined from independent triplicate curve fits, 18 data points per curve)
AOC = Area Over Curve
Day = Age of culture after 72h drug treatment
reduction or gain in drug sensitivity ($IC_{50}$ or $GR_{50}$ 2D vs 3D value changes > 5x, AOC value changes > 1.5x)

$^{*}$ $p < 0.05$, $^{**}$ $p < 0.01$

### Supplementary Table S4 | Summary of 2D vs 3D drug-sensitivity metrics for MiaPaCa-2.

MiaPaCa-2 cells were cultured in 2D (day 4) or in an ECM-containing 3D hydrogel (day 7 or day 11) and treated for 72 h with the indicated agents. Columns report $IC_{50}$, $GR_{50}$, and GR_AOC (area over the GR curve), derived from independent triplicate dose–response fits (18 data points per curve). Values reported as “>10” or “>100” indicate that the $IC_{50}$ or $GR_{50}$ exceeded the highest concentration tested. Lower $IC_{50}$/$GR_{50}$ values and lower GR_AOC indicate higher drug sensitivity. Color shading encodes the direction and magnitude of change relative to 2D day 4 (pink, decreased sensitivity in 3D; green, increased sensitivity in 3D; thresholds: ≥5× change for $IC_{50}$/$GR_{50}$ and ≥1.5× for GR_AOC). Asterisks denote significant differences between 3D and 2D conditions (*$P < 0.05$; **$P < 0.01$).

**Overall activity summary:** Three of five standard-of-care agents (9/14 total drugs tested) retain activity at 3D day 7 ($P < 0.05$), whereas none of the five standard-of-care agents (2/14 total drugs tested) remain active after extended 3D growth (day 11; $P < 0.05$). Consistent with GR dose–response curves, MiaPaCa-2 cells exhibit progressive loss of drug sensitivity with prolonged 3D residence, most evident by day 11 for 5-FU, gemcitabine, and paclitaxel, with occasional transient gains at day 7 for selected targeted agents that are not maintained at day 11.

4 out of 5 standard of care drugs (13 of 14 total test drugs) remain active after 7 days 3D growth ($p < 0.05$)
4 out of 5 standard of care drugs (13 of 14 total test drugs) remain active after extended 3D growth ($p < 0.05$)

| | CFPAC-1 2D $IC_{50}$ Day4 | CFPAC-1 3D $IC_{50}$ Day7 | CFPAC-1 3D $IC_{50}$ Day11 | CFPAC-1 2D $GR_{50}$ Day4 | CFPAC-1 3D $GR_{50}$ Day7 | CFPAC-1 3D $GR_{50}$ Day11 | CFPAC-1 2D GR_AOC Day4 | CFPAC-1 3D GR_AOC Day7 | CFPAC-1 3D GR_AOC Day11 |
|---|---|---|---|---|---|---|---|---|---|
| 5-FU | 5.57 | 7.86 | NF | 4.32 | 0.387 | 13.8 | 1.33 | 2.25 | 0.44 |
| GEM | 0.00633 | 0.0128 | 2.36 | 0.00534 | 0.00790 | 0.0179 | 3.73 | 3.32 | 3.12 |
| OXA | > 10 | > 10 | NF | 14.7 | NF | NF | 0.06 | 0.92 | -1.04 |
| PAC | 2.32E-07 | NF | 62.4 | 2.01E-09 | 0.00149 | 0.0366 | 6.64 | 3.18 * | 2.45 * |
| SN38 | 0.00228 | 0.00123 | 0.00345 | 0.00175 | 0.000681 | 0.000474 | 4.34 | 4.78 | 6.53 ** |
| BOR | 0.00404 | 0.00425 | 0.00541 | 0.00376 | 0.00255 | 0.00278 | 3.92 | 3.87 | 6.70 ** |
| CAR | 10.6 | > 10 | 5.81 | 9.25 | NF | 3.09 | 0.29 | 0.85 | -0.19 |
| DIS | > 10 | > 10 | NF | NF | NF | 5.19 | 0.10 | 0.24 | -0.29 |
| DOX | 0.471 | 0.349 | 1.45 | 0.401 | 0.104 | 0.113 | 1.56 | 2.21 | 2.88 * |
| HOM | 0.0592 | 0.0444 | 0.111 | 0.0556 | 0.0264 | 0.0299 | 2.99 | 2.58 | 4.03 |
| KPT | 0.0606 | 0.0640 | 2.50 | 0.0479 | 0.0260 | 0.0119 | 2.61 | 2.44 | 3.37 |
| OBA | 0.240 | 0.263 | 2.29 | 0.216 | 0.171 | 0.275 | 1.94 | 1.38 | 1.22 |
| TRA | 60.3 | NF | 0.537 | 18.3 | 0.00585 | 0.0000136 | 0.89 | 3.04 | 4.94 * |
| YM155 | 0.00465 | 0.00443 | 0.00615 | 0.00438 | 0.00371 | 0.00230 | 3.70 | 3.55 | 6.71 ** |

$IC_{50}$ / $GR_{50}$ values in uM (determined from independent triplicate curve fits, 18 data points per curve)
AOC = Area Over Curve
Day = Age of culture after 72h drug treatment
reduction or gain in drug sensitivity ($IC_{50}$ or $GR_{50}$ 2D vs 3D value changes > 5x, AOC value changes > 1.5x)

* $p < 0.05$, ** $p < 0.01$

### Supplementary Table S5 | Summary of 2D vs 3D drug-sensitivity metrics for CFPAC-1.

CFPAC-1 cells were cultured in 2D (day 4) or in an ECM-containing 3D hydrogel (day 7 or day 11) and treated for 72 h with the indicated agents. Columns report $IC_{50}$ (µM), $GR_{50}$ (µM), and GR_AOC (area over the GR curve; unitless), derived from independent triplicate dose–response fits (18 data points per curve). Values shown as ">10" or ">100" exceeded the highest concentration tested. Lower $IC_{50}$/$GR_{50}$ and lower GR_AOC indicate higher sensitivity. Color shading encodes the direction and magnitude of change relative to 2D day 4 (pink, decreased sensitivity in 3D; green, increased sensitivity in 3D; thresholds: ≥5× change for $IC_{50}$/$GR_{50}$ and ≥1.5× for GR_AOC). Asterisks denote significant differences between 3D and 2D conditions (*$P < 0.05$; **$P < 0.01$).

**Overall activity summary:** Four of five standard-of-care agents (13/14 total drugs tested) retain activity at 3D day 7 ($P < 0.05$), and four of five remain active after extended 3D growth (day 11; $P < 0.05$). Consistent with GR dose–response curves, CFPAC-1 exhibits multiple instances of increased or sustained drug sensitivity in 3D, including durable enhancement for selected agents between day 7 and day 11 (e.g., SN-38 and trametinib).

### Supplementary Table S6 | Comparison of $GR_{50}$ values with clinical plasma exposure across PDAC cell lines and culture formats.

| | MiaPaca2 | MiaPaca2 | CFPAC-1 | CFPAC-1 | Panc-1 | Panc-1 |
|---|---|---|---|---|---|---|
| | 2D GR50 Day4 | 3D GR50 Day11 | 2D GR50 Day4 | 3D GR50 Day11 | 2D GR50 Day4 | 3D GR50 Day11 |
| 5-FU | 4.13 | > 100 | 4.32 | 13.8 | 6.39 | 105 |
| GEM | 0.00743 | > 10 | 0.00534 | 0.0179 | NF | 30.1 |
| OXA | 1.45 | > 10 | 14.7 | NF | 0.592 | 4.72 |
| PAC | 5.96E-07 | > 10 | 2.01E-09 | 0.0366 | 1.57E-07 | 0.186 |
| SN38 | 0.0282 | 0.387 | 0.00175 | 0.000474 | 0.0921 | 0.367 |

| | | | MiaPaca2 | MiaPaca2 | CFPAC-1 | CFPAC-1 | Panc-1 | Panc-1 |
|---|---|---|---|---|---|---|---|---|
| | Mean Cmax (µM) | Mean t1/2 | 2D GR50 Day4 | 3D GR50 Day11 | 2D GR50 Day4 | 3D GR50 Day11 | 2D GR50 Day4 | 3D GR50 Day11 |
| 5FU20 | 426 | 0.32 hours | ~100x | ≤ 4x | ~100x | ~30x | ~70x | ~4x |
| GEM21 | 5.05 | 0.3 hours | ~700x | ≤ 0.5x | ~1000x | ~300x | - | ~0.2x |
| OXA22 | 3.6 | 0.24 hours | ~2.5x | ≤ 0.4x | ~0.25x | - | ~6x | ~0.8x |
| PAC23 | 6.9 | 8 - 34 hours | >106x | ≤ 0.7x | >106x | ~200x | >106x | ~40x |
| SN3824 | 0.11 | 12.8 hours | ~4x | ≤ 0.3x | ~60x | ~200x | ~1x | ~0.3x |

**(Upper table)** Mean $GR_{50}$ values (µM) for five standard-of-care agents—5-fluorouracil (5-FU), gemcitabine (GEM), oxaliplatin (OXA), paclitaxel (PAC), and SN-38—measured in MiaPaCa-2, CFPAC-1, and PANC-1 cells under 2D (day 4) and extended 3D (day 11) culture conditions. Values were derived from growth-rate–normalized 72-h dose–response curves. Shaded cells highlight drug–cell line combinations where extended-3D $GR_{50}$ values approach or exceed reported physiological plasma concentrations.

**(Lower table)** Reported mean patient plasma maximum concentrations (Cmax) and elimination half-lives ($t_{1/2}$) for the same agents, compiled from clinical pharmacokinetic studies (refs. 20–24). Right-hand columns indicate the approximate fold-difference between clinical Cmax and corresponding in-vitro $GR_{50}$ values (e.g., “~100×” indicates that plasma levels exceed $GR_{50}$ by two orders of magnitude). While clinical Cmax values are typically 1–2 orders of magnitude higher than 2D $GR_{50}$ values, they are often comparable to or lower than extended-3D $GR_{50}$ values, particularly in MiaPaCa-2 and PANC-1, indicating that prolonged 3D culture reveals drug-tolerance levels more consistent with clinically achievable exposures.

### Supplementary Table S7 | Influence of spheroid size on microenvironmental organization and therapeutic response.

| Spheroid Size | Characteristics | Comparison to 2D Cultures | Pathophysiological Features | In Vivo Analogy |
|---|---|---|---|---|
| **<150 µm** | 3D cell–cell and cell–matrix interactions<br>Altered expression profile | Different proliferative activity than 2D cultures | Limited or not yet established<br>No inward/outward pathophysiological gradients yet | Early tumor aggregates, before strong gradients form |
| **200–500 µm** | Development of chemical gradients (oxygen, nutrients, catabolites) | More complex than 2D cultures, start to mimic tissue microenvironment | Establishment of radial proliferative and metabolic gradients | Comparable to small avascular tumor nodules or micro metastases |
| **>500 µm** | Central hypoxia and necrosis may form | Strong divergence from 2D due to heterogeneous zones | Peripheral cells actively cycling (like near capillaries)<br>Inner cells quiescent, dying via apoptosis/necrosis<br>Clear concentric heterogeneity | Reflects inter-capillary microregions of solid tumors |

Summary of characteristic biological and pathophysiological features associated with increasing spheroid size in 3D tumor models. As spheroids grow, progressive establishment of diffusion-limited gradients (e.g., oxygen, nutrients, metabolites) leads to spatial heterogeneity in proliferation, metabolism, and viability, resulting in treatment responses that increasingly diverge from 2D monolayer cultures. Approximate in vivo analogies are provided to contextualize spheroid size–dependent microenvironmental states observed in solid tumors. Adapted from Hirschhaeuser et al., 2010.

---

*Hirschhaeuser, F., Menne, H., Dittfeld, C., West, J., Mueller-Klieser, W., and Kunz- Schughart, L. A. (2010).*
***Multicellular Tumor Spheroids: An Underestimated Tool Is Catching up Again*.**
*J. Biotechnol. 148, 3–15. doi:10.1016/ j.jbiotec.2010.01.012*

### Supplementary Table S8 | Compounds used in this study and their reported mechanisms of action.

| Compound | Abbreviation | Mechanism(s) of action | Reported ABC transporter interaction* | Example primary reference(s)** |
|---|---|---|---|---|
| KPT-185 | KPT | CRM1/XPO1 inhibitor; induction of apoptosis | ABC substrate status not established | Etchin 2013, Blood (KPT-330/selinexor) |
| Obatoclax | OBA | Pan-BCL-2 family inhibitor; autophagy modulation | Not an ABCB1 substrate; reported inhibitor of ABCB1/ABCG2; MDR-reversing agent | Qiu 2012, Biochem Pharmacol; Xiao 2021 |
| Trametinib | TRA | MEK1/MEK2 inhibitor | Context-dependent interaction with ABCB1; reported to modulate ABCB1-mediated MDR, not a classic substrate | Qiu 2015, Oncotarget |
| YM-155 | YM | Survivin suppressor | Substrate of ABCB1 and ABCG2 | Imai 2012, Cancer Sci; Xiao 2021 |
| Idarubicin | IDA | Topoisomerase II inhibitor;<br>DNA damage response; autophagy | Substrate of ABCB1; efflux by ABCC1 and other MRPs | Szakács 2006, Nat Rev Drug Discov; Xiao 2021 |
| Homoharringtonine | HOM | Protein translation inhibitor | ABC transporter interactions not well defined | Chen 2009, Leukemia |
| Disulfiram | DIS | Aldehyde dehydrogenase inhibitor; ROS and metal-dependent cytotoxicity | Not a direct ABC substrate; reported to modulate MDR indirectly (e.g., via ALDH/ROS/metal complexes) | Triscott 2012, Oncotarget; Xiao 2021 |
| Bortezomib | BOR | Proteasome inhibitor; autophagy induction | Not a primary substrate of ABCB1/ABCG2; limited direct efflux | Minderman 2004, Clin Cancer Res; Xiao 2021 |
| 5-Fluorouracil | 5-FU | Antimetabolite; thymidylate synthase inhibition;<br>DNA/RNA synthesis disruption | Not a substrate of major ABC transporters (ABCB1, ABCC1, ABCG2) | Longley 2003, Nat Rev Cancer |
| Gemcitabine | GEM | Nucleoside analog; DNA synthesis inhibition;<br>replication fork stalling | Efflux of gemcitabine metabolites by ABCC (MRP) family (e.g., ABCC4/ABCC5) | Oguri 2007, Cancer Sci; Xiao 2021 |
| Oxaliplatin | OXA | Platinum-based DNA crosslinking agent;<br>DNA damage response | Efflux by ABCC2 and other MRPs reported; contributes to platinum resistance | Burger 2010, Pharmacol Ther; Xiao 2021 |
| Paclitaxel | PAC | Microtubule stabilizer; mitotic arrest | Canonical ABCB1 (P-gp) substrate | Szakács 2006, Nat Rev Drug Discov; Xiao 2021 |
| SN-38 | SN-38 | Topoisomerase I inhibitor; DNA damage response | Substrate of ABCG2 and ABCB1 | Nakatomi 2001, Int J Cancer; Xiao 2021 |
| Carboplatin | CAR | Platinum-based DNA crosslinking agent;<br>DNA damage response | Efflux by ABCC (MRP) family reported; generally weaker than cisplatin/oxaliplatin | Burger 2010, Pharmacol Ther; Xiao 2021 |
| Doxorubicin | DOX | Topoisomerase II inhibitor;<br>DNA intercalation; oxidative stress | Canonical substrate of ABCB1; also transported by ABCC1 and other MRPs | Szakács 2006, Nat Rev Drug Discov; Xiao 2021 |

Table lists investigational and approved compounds tested in 2D and 3D culture formats, along with abbreviations used throughout the manuscript and their primary reported mechanisms of action based on published literature.

---

** ABC transporter associations are based on reported substrate or efflux relationships in cancer models and clinical literature; strength and relevance may vary depending on cellular context, transporter expression, and microenvironmental conditions.*
